# CD81 partners with CD44 in promoting exosome biogenesis, tumor cluster formation, and lung metastasis in triple negative breast cancer

**DOI:** 10.1101/2022.02.23.481674

**Authors:** Erika K. Ramos, Chia-Feng Tsai, Nurmaa K. Dashzeveg, Yuzhi Jia, Yue Cao, Megan Manu, Rokana Taftaf, Andrew D. Hoffmann, Lamiaa El-Shennawy, Marina A. Gritsenko, Valery Adorno-Cruz, Emma J. Schuster, David Scholten, Dhwani Patel, Xia Liu, Priyam Patel, Brian Wray, Youbin Zhang, Shanshan Zhang, Ronald J. Moore, Jeremy V. Mathews, Matthew J. Schipma, Tao Liu, Valerie L. Tokars, Massimo Cristofanilli, Tujin Shi, Yang Shen, Huiping Liu

## Abstract

Tumor-initiating cells with reprogramming plasticity are thought to be essential for cancer development and metastatic regeneration in many cancers; however, the molecular mechanisms are not fully understood. This study reports that CD81, a tetraspanin protein marker of small extracellular vesicles (exosomes), functions as a binding partner of CD44 and facilitates self-renewal of tumor initiating cells. Using machine learning-assisted protein structure modeling, co-immunoprecipitation, and mutagenesis approaches, we further demonstrate that CD81 interacts with CD44 on the cellular membrane through their extracellular regions. In-depth global and phosphoproteomic analyses of clustering tumor cells unveils endocytosis-related signature pathways of proteins and phosphorylation patterns regulated by CD81 and CD44 specifically or shared between two. Notably, CRISPR Cas9-mediated depletion of either CD44 or CD81 results in loss of both proteins in cancer cell-secreted exosomes, a state which abolishes exosome-induced self-renewal of recipient cells for mammosphere formation. CD81 is expressed in >80% of human circulating tumor cells (CTCs) and specifically enriched in clustered CTCs along with CD44 isolated from breast cancer patients. Mimicking the phenotypes of CD44 deficiency, loss of CD81 also inhibits tumor cluster aggregation, tumorigenesis, and lung metastasis of triple negative breast cancer (TNBC), supporting the clinical significance of CD81 in association with patient outcomes. Our study highlights the novel role of CD81 and its partnership with CD44 in cancer exosomes, self-renewal, CTC clustering, and metastasis initiation of TNBC.

## Introduction

Breast cancer killed nearly 0.7 million people worldwide in 2020 with metastasis accounting for 90% of deaths (1). Negative for estrogen receptor, progesterone receptor, and HER2 amplification, triple negative breast cancer (TNBC) comprises 10-15% of newly diagnosed breast cancer cases and is highly metastatic with low long-term survival (2–5). TNBC preferentially metastasizes to the visceral organs such as the lungs, liver, and brain (6, 7). Growing evidence suggests tumor-initiating cells, or cancer stem cells with self-renewal and regenerative reprogramming capacity, are the underlying cause of cancer relapse, therapy resistance, and distant dissemination (8–20) with measurable clinical impact on patient outcomes (16, 21–31). However, the cellular and molecular mechanisms contributing to tumor cell spreading and regenerative metastasis are yet to be fully understood.

Circulating tumor cells (CTCs) with genetic and/or epigenetic alterations (32, 33) are tumor cells that shed from cancer lesions and circulate within the blood or lymphatic vasculature before seeding with a very low frequency to distant organs for metastatic tumor regeneration. Detection of CTC events on the CellSearch platform is associated with patient outcomes in which multicellular CTC clusters predict worse prognosis and mediate metastasis in a 20- to 100-fold higher efficiency than single CTCs (34–36). Our recent work demonstrated that the breast tumor initiating cell marker CD44 is expressed in around 80% of CTC clusters and predicts an unfavorable overall survival of patients with breast cancer, especially TNBC (36). CD44 mediates homophilic interactions for CTC cluster formation as a mechanism to promote self-renewal (mammosphere formation) and polyclonal metastasis in TNBC (36). While most of the previous studies focus on the genome, epigenome, or transcriptome alterations in clustering tumor cells (33, 35, 37), little is directly linked to proteome alterations. We performed mass spectrometry proteomic profiling of TNBC patient-derived xenografts (PDX) tumor cells that cluster and discovered a tetraspanin protein and exosome marker CD81 as one of the altered proteins upon CD44 depletion. The objective of the study is to elucidate the functions of CD81 in self-renewal and progression of TNBC, and to explore its potential connection with cancer exosomes, all of which are extremely understudied.

Exosomes are cell-secreted extracellular vesicles (EVs) (30-140 nm) contributing to pivotal intercellular communicator and serving as a new biologic nanoparticle platform for therapeutic development and targeted drug delivery (38–45). Distinct from the biogenesis pathway of large EVs (such as microvesicles) *via* plasma membrane shedding, exosomes are small EVs generated from the invagination of the plasma membrane to form endosomes that continue to invaginate and mature into multivesicular bodies destined for either lysosome degradation or fusion with the plasma membrane and release as exosomes (46). Exosomes are characterized by presence of membrane structure and enriched protein markers such as CD81, CD63, CD9, and TSG101 (46, 47). Exosomes have emerged as key players in cancer development (48, 49) and distant organ-specific metastasis (50); however, the role of exosome markers such as CD81 in tumor initiation and progression remains relatively unclear. This work aims to determine the role of CD81 in cancer exosomes, tumor initiation, and metastasis. Following the widely adopted nomenclature in the EV field (51) and considering the technical limitations in exosome isolation that contains heterogenous EV populations, we therefore utilize “EVs” to describe the enriched exosomes in our study.

Here, using the TNBC PDXs established and maintained in mice (16, 36), human cell lines, and mouse tumor models with genetic modulations such as knockdown (KD) and knockout (KO), we examined the importance of CD81 in TNBC progression and identified a novel mechanistic link to CD81/CD44 interactions and self-renewal associated with cancer exosomes, and identified previously unknown tumor-intrinsic functions of CD81 in promoting tumor clustering and lung metastasis.

## Results

### CD81 promotes mammosphere formation and interacts with CD44

We previously found that TNBC PDXs express splicing variants of CD44 (CD44v), whereas MDA-MB-231 cells predominantly express standard CD44 (CD44s), both contributing to CTC cluster formation and cancer metastasis. To better understand the proteomic alterations and mechanistic regulation of self-renewal and tumor cluster formation, we conducted global mass spectrometry analyses of TNBC cells, including clustering and non-clustering PDX tumor cells (CD44^+^ and CD44^−^ cells) as well as MDA-MB-231 cells of wild-type (WT) and *CD44* KO, which pooled populations were generated via multiple guide RNAs and CRISPR-Cas9 technique (36). Based on two paired comparisons with PDX cells freshly isolated from mice, including flow-sorted CD44^+^ versus CD44^−^ cells (mimicking circulation) and siCD44 mediated KD versus siRNA control cells that clustered *ex vivo* (36), we identified 38 overlapping proteins differentially expressed in these cells with a more than 2-fold change (**Supplementary Table S1**). In addition to PAK2 as a previously reported CD44 target (36), one of the top listed proteins was CD81, an exosome marker and tetraspanin family membrane protein, reduced in sorted CD44^−^ or CD44 KD cells. Consistently, CD81 was present in the TNBC PDX (CD44v) cells but decreased in CD44KO PDXs (**Supplementary Figure S1A**). However, in MDA-MB-231 cells that express CD44s, CD81 was only temporarily down-regulated in CD44KO cells in suspension but comparable between adherent CD44 WT and KO cells, as detected by immunoblotting (**Supplementary Figure S1B**), suggesting context dependent alterations of CD81 by CD44 in TNBC cells.

Since CD44 is known to promote tumor initiation and metastasis in breast cancer (36), we first tested the role of CD81 in self-renewal in TNBC by assessing its impact on mammosphere formation, cell growth, and pluripotency-related genes and proteins after gene KO, which was generated as pooled populations with two CD81 gRNAs using the CRISPR-Cas9 approach (**Supplementary Figure S1B**). In addition to CD44KO and CD81KO, a combined CD44/CD81 double KO (dKO) cells were also made for phenotypic analyses (**Supplementary Figure S1B**). In comparison to the WT MDA-MB-231 cells, pooled populations of CD44KO, CD81KO, and dKO cells show similar, diminished capability for mammosphere formation, with fewer and smaller spheres where the dKO did not show much additional effects than single KOs (**Figure 1A, B**), suggesting both CD81 and CD44 are required for optimal self-renewal with similar functional importance.

**Figure 1.**
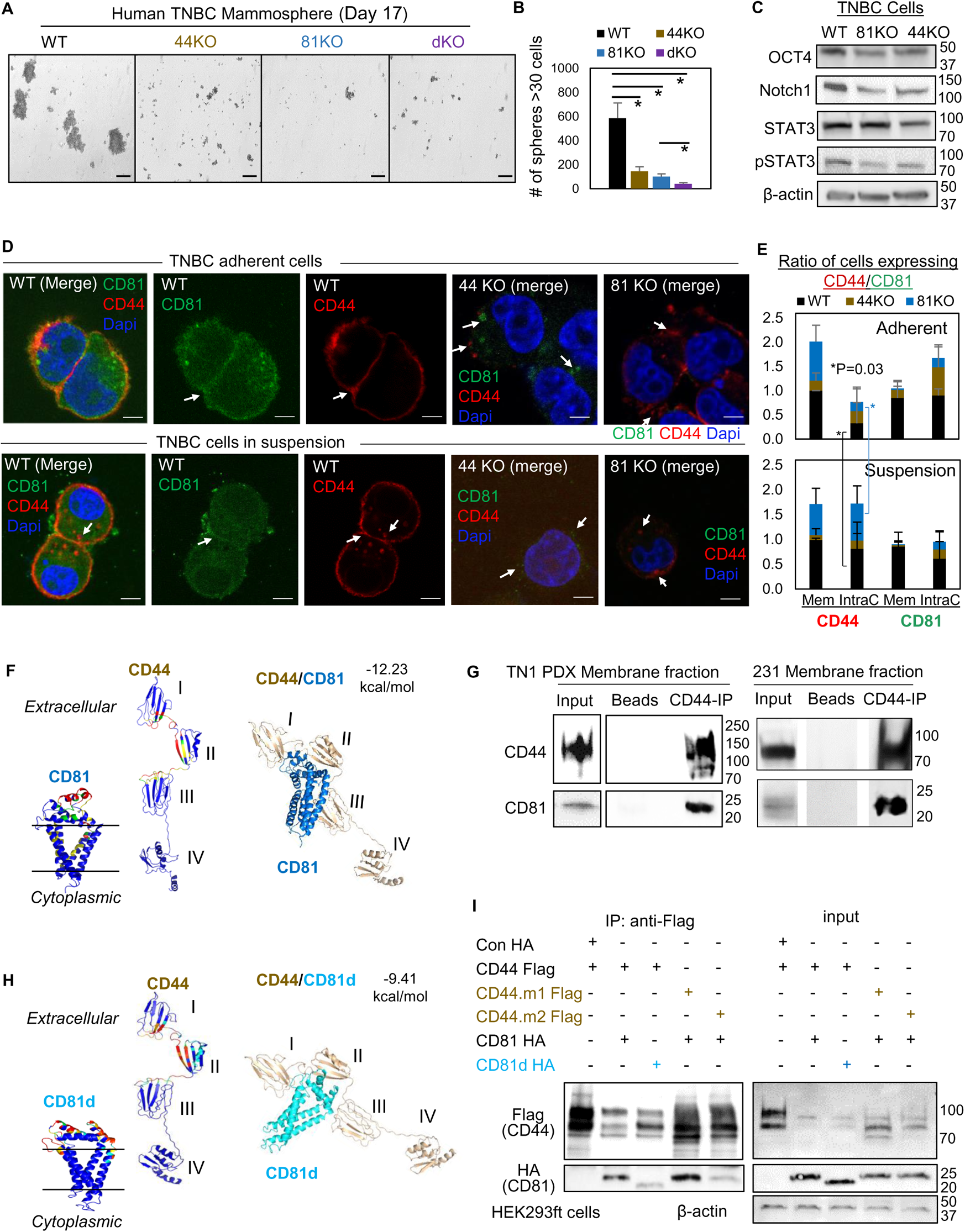
CD81 interacts with CD44 on the membrane and promotes mammosphere formation of TNBC cells. **A, B.** Representative images (A) and bar graphs (B) of the mammospheres of MDA-MB-231 cell groups (WT, CD44KO, CD81KO, dKO pool populations), 17 days after seeded at 2,000 cells per well (6-well plate) in serum-free mammosphere formation media. N=4 replicates. Repeated 3 times. Scale bar = 100 µm. One-tailed student T-test * P < 0.05. **C.** Immunoblot analysis of OCT4, Notch1, STAT3, and pSTAT3 expression in WT, CD81KO, and CD44KO MDA-MB-231 cells (pooled KO cells). **D.** Immunofluorescence of CD44 and CD81 in adherent or in suspension MD-MB-231 cells (WT, CD44KO, and CD81KO) stained with anti-CD44-Texas Red, anti-CD81-Alexa488, and DAPI for DNA. Examined 20 WT, 14 CD44KO, and 20 CD81KO adherent cells and 31 WT, 31 CD44KO, and 28 CD81KO suspension cells. Scale bar =5 µm. **E.** Bar graphs of membrane and intracellular CD44 and CD81 localization in WT, CD44KO, and CD81KO MDA-MB-231 cells. Anova analyses among three groups with P values for adherent and in-suspension cells, respectively: 0.0004 and 7.357E-08 (membrane CD44), 0.9119 and 0.0024 (intracellular CD44), 0.0022 and 1.079E-09 (membrane CD81), and 0.0255 and 0.0103 (intracellular CD81). Two-tailed student T-test P= 0.03 for intracellular CD44 levels between adherent and suspension cells (both WT and CD81KO cells). **F.** Predictive modeling of CD44 and CD81 interaction with hot spots shown in red and yellow. Top-ranked structural models of predictive interactions between CD44/CD81 (estimated binding energy: −12.23 Kcal/mol) and between CD44/CD81d (deletion mutant) (estimated binding energy: −9.41 Kcal/mol). **G.** Immunoblots of endogenous CD44 and CD81 immunoprecipitated by anti-CD44 from the lysates (membrane fraction) of MDA-MB-231 and (total) TN1 PDX cells. **H.** Predictive modeling of CD44 and CD81d (deletion mutant) interaction with hot spots shown in red and yellow. Top-ranked structural models of predictive interactions between CD44/CD81d (deletion mutant) (estimated binding energy: −9.41 Kcal/mol). **I.** Representative immunoblots of CD81-HA (CD81d) immunoprecipitated by anti-CD44-Flag (CD44 mutants CD44.1 and CD44.2) from the lysates of HEK293ft cells (N=3 biological replicates; Con, control). Cells were transfected with either Flag-CD44 or mutants (Flag-CD44.1, Flag-CD44.2) and Con HA, HA-CD81, or HA-CD81d with deletion at amino acids 159-187 (CD81d). 48 hours after transfection the cells were harvested for Flag IP.

Using multiple mouse Cd81 gRNAs and CRISPR-Cas9 approach, we also created pooled Cd81KO populations in mouse 4T1 TNBC cells which displayed fewer mammospheres compared to the WT cells, after being seeded with a low number of cells in stem cell medium in suspension, (**Supplementary Figure S1C**). To assess human CD81 and mouse Cd81 functions in cell growth, we employed IncuCyte time-relapse imaging to monitor the cell confluence in adherent culture. Both CD81KO and Cd81KO cells showed a slightly less confluency compared to respective WT cells (**Supplementary Fig S1D-E**), indicating that effects of CD81KO on diminished self-renewal (mammosphere formation) are beyond a slightly altered cell growth or proliferation.

We then measured the expression levels of stem-cell signature genes and/or proteins in these cells. Like CD44KO cells, CD81 KO populations decreased protein levels of breast tumor-initiating markers including OCT4, NOTCH1, and phosphoSTAT3 (**Figure 1C**). In the meantime, siRNA-mediated transient KD of CD81 slightly reduced mRNA expression of the genes *CD44*, *CD47* and *NF-kB* (**Supplementary Figure S1F**).

To determine whether the two membrane proteins CD44 and CD81 influence each other’s cellular localization or distribution, we further performed immunofluorescence staining of human TNBC cells with CD44KO or CD81KO, adherent and in suspension (for 3 h clustering). CD44 was observed mostly on the cytoplastic membrane in adherent WT MDA-MB-231 cells but drastically accumulated to the intracellular regions in suspension cells (P=0.03) (**Figure 1D-E**). In the WT group, ~50% of adherent cells and 70% of suspension cells showed an average of 14-16% of partial colocalization between CD44 and CD81 on the cytoplasmic membrane, with CD81 mainly presented at the interface for clustering tumor cells in suspension (**Figure 1D, Supplementary Figure S1G**). The KO of CD44 or CD81 altered the localization of CD81 or CD44, respectively, with disrupted localization to the membrane but increased staining at intracellular loci of adherent cells (**Figure 1D**, white arrows in top panels). Meanwhile, CD44KO or CD81KO further weakened the detection of both in either intracellular loci or surface membrane in suspension cells (**Figure 1D**, white arrows in bottom panels). These data demonstrate that cell detachment influences the cellular localization of CD81 and CD44 which facilitate each other’s membrane presentation and subsequent co-localization, especially at the interface of neighboring and clustering tumor cells.

### CD81 forms a protein complex with CD44 dependent on extracellular regions

To further examine if CD81 directly interacts with CD44 and to identify possible molecular regions responsible for proposed CD44-CD81 interactions in TNBC cells, we employed machine learning-assisted protein structure modeling (52) and co-immunoprecipitation (co-IP) tests. Based on previous structural studies on CD44 (53) and CD81 (54) as well as computational programs iTasser (55), ClusPro (56), and Bayesian Active Learning (BAL) (52), we first analyzed CD44 and CD81 protein sequences and possible dynamics of protein-protein interaction models. As shown in **Figure 1F**, the hotspot residues in warm colors (red and yellow) predicted to be involved in the interactions are located in domains I and II of CD44, which also contribute to CD44-CD44 homophilic interactions (36, 53), and the extracellular loop of CD81, which links its third and fourth transmembrane domains, with an estimated free energy of binding at −12.23 kcal/mol. As expected, after being immunoprecipitated (IP) by anti-CD44 antibody, the endogenous CD81 and CD44 proteins were simultaneously observed within a membrane-fraction protein complex from the lysates of TN1 PDXs (expressing CD44v) and MDA-MB-231 cells (expressing CD44s) (**Figure 1G**). We then overexpressed tagged CD44-Flag and CD81-HA in CD44^−^ HEK293T cells, in which CD44 was shown in multiple forms at distinct molecular weights (**Supplementary Figure S1H**), possibly due to variable glycosylation patterns as we previously reported (53). Both CD44-Flag and CD81-HA were detected in the protein complex after co-IP with anti-Flag magnetic beads (**Supplementary Figure S1H**), demonstrating the interaction of exogenous CD44 and CD81 in these cells.

According to predicted protein structures for CD81 and CD44 interactions, we further designed a deletion mutant of CD81d by removing the loop region of amino acids 159-187 (CD81d), which expression was stable and presented on the cell membrane (**Supplementary Figure S1I-J**), and predicted to dramatically impair the interaction, with an estimated free energy of binding altered to −9.41 kcal/mol (**Figure 1H**). We then assessed the effects of CD81d on its interaction with CD44, and the effects of CD44 mutants CD44.m1 and CD44.m2, with point mutations in domains I and II (53), respectively, on the interaction with CD81 (**Supplementary Figure S1I**). By three independent co-IP tests using the tagged CD44-Flag and CD81-HA co-transfected cells, we found that CD81d and CD44.m2 mutant partially interfered with the CD44-CD81 interactions (**Figure 1I**), indicating the specific regions required for CD81-CD44 interactions.

### Shared and distinct pathways regulated by CD81 and CD44 in self-renewal, proliferation, and endocytosis

To elucidate the membrane protein CD81-regulated molecular networks and pathways in self-renewal, we analyzed mass spectrometry-based global proteomes and phosphoproteomes as well as RNA sequencing-based transcriptomes of TNBC cells (adherent or in suspension) with CD81 and CD44 depletion.

We first performed RNA sequencing to examine the CD81 KD effects on transcriptome in adherent MDA-MB-231 cells after siCD81 transfections (**Supplementary Figure S2A**). The Metascape pathway (57) analysis identified CD81 KD-influenced transcriptome in the pathways of protein modification, cell proliferation, and differentiation (**Supplementary Figure S2B**). However, among a few hundreds of significantly altered transcripts, most of them had very low baseline detection, and only a few genes with robust expression were up-regulated over 2-fold (such as *SEMA7a*, *HMGA2*, *ACVR2B*, *YOD1*, and *FUT4*) or down-regulated more than half, including *CDC34*, *UBE2R2*, *SLC7A11*, and *ADIRF* (**Supplementary Figure S2C, Supplementary Table S2, Excel Data 1**). Among the siCD81-upregulated genes, SEMA7a, a glycosylphosphatidylinositol membrane anchor promoting osteoclast and blood cell differentiation (58, 59), when further depleted in CD81KO cells, siSEMA7a partially rescued or restored mammosphere formation in these cells (**Supplementary Figure S2D-F**), suggesting a role of SEMA7a in inhibiting self-renewal of breast cancer cells.

To further explore the possible proteome alterations connecting CD81 with CD44 regulation in TNBC cells, we pursued mass spectrometry analyses of these cells with transient KDs. Our pilot proteomic comparisons between adherent versus clustering MDA-MB-231 cells in-suspension, showed minimal protein level alterations (5 proteins with >2-fold changes) but drastic phosphoproteomic alterations (over 1,300 phosphopeptides with >2-fold changes) (**Supplementary Figure S3A, Excel Data 2**), suggesting that posttranslational phosphorylation significantly modulates the signaling pathways of cells in suspension that mimick detached migrating cells and CTCs.

We then collected three sets of control, siCD81 and siCD44-transfected cells in suspension (3 h clustering) in which a specific depletion of CD81 or CD44 was achieved (**Supplementary Figure S3B**) for both global proteome and phosphoproteome analyses. By comparative mass spectrometry analyses of three groups of cell lysates, we identified 6 clusters of altered general proteins (G1-G6) and another 6 clusters of altered phosphoproteins (P1-P6) with top pathways for each cluster annotated by KEGG (60) (**Figure 2A-D, Supplementary Tables S3, Excel Data 3-4**). In comparison to the control cell profiles, Clusters G1 (downregulated) and G6 (upregulated) represent the altered proteins in both siCD81 and siCD44 cells in the same directions, suggesting shared pathway regulation, such as DNA replication and cell cycle in G1 and metabolic pathways in G6, involved in the regulation of self-renewal and cell proliferation. Ribosome pathway was the top pathway in Clusters G3 and G4 with downregulated proteins specific to siCD44 KD and siCD81 KD, respectively, suggesting distinct pathway components but possible contribution to similar ribosome-regulating functions of CD81and CD44 in TNBC cells.

**Figure 2.**
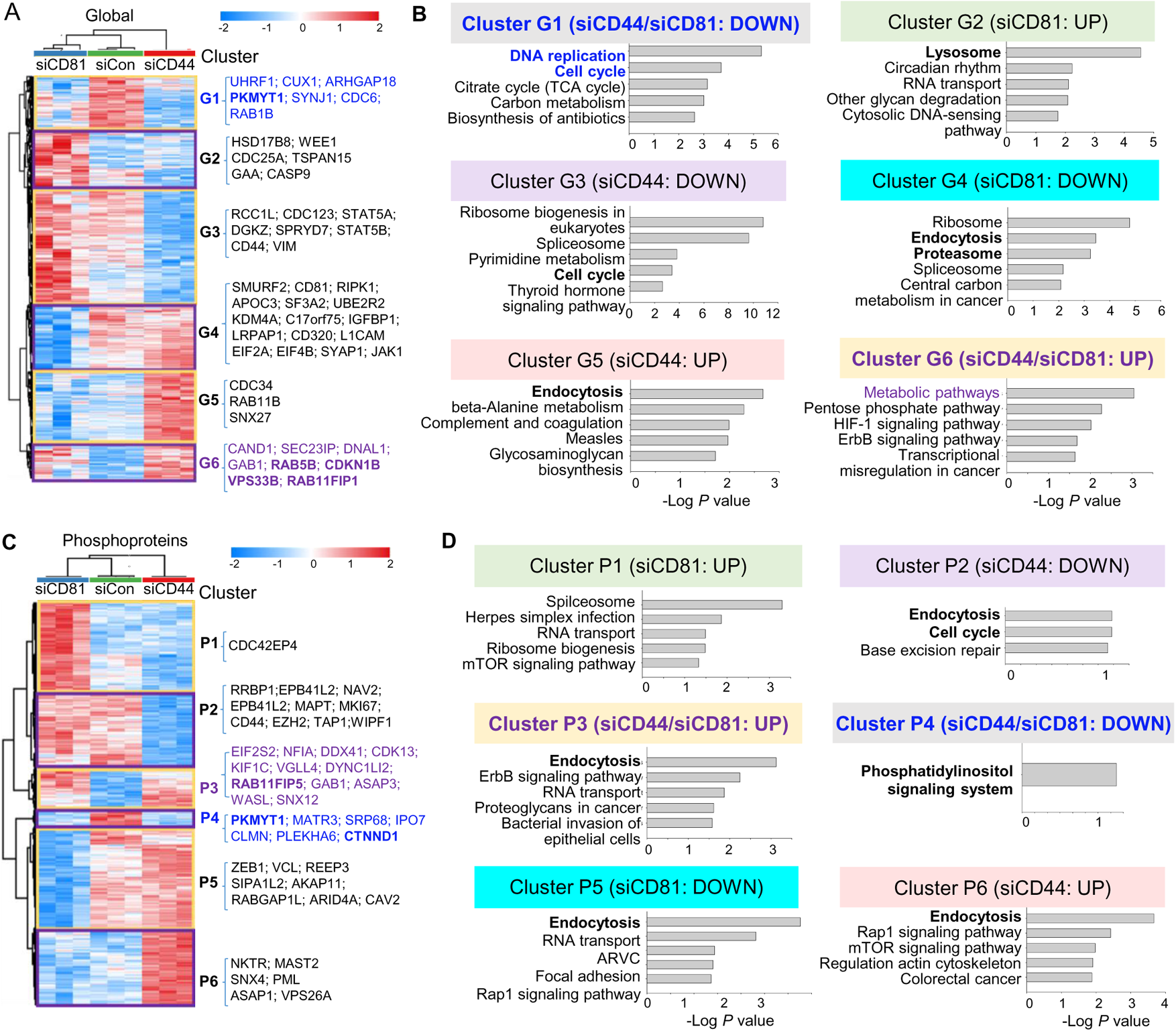
Global mass spectrometry and phosphoproteomic profiling of MDA-MB-231 cells with siCD81 and siCD44 KDs. **A, B.** Global mass spectrometry heatmap (A) and KEGG pathway analysis (B) of altered top pathways with significantly expressed proteome within 6 different clusters (G1-G6) in siControl, siCD81, and siCD44 cells at 3 h in suspension (N=3 replicates, Anova t-test FDR<0.01, P=<0.05)). **C, D.** Phosphoproteomic mass spectrometry heatmap (C) and KEGG pathway analysis (D) of altered top pathways with significantly changed phosphoproteome within 6 different clusters (P1-P6) in siControl, siCD81, and siCD44 cells at 3 h in suspension (N=3 replicates, Anova t-test FDR<0.01, P=<0.05)

Most notably, endocytosis, lysosome, and proteosome pathways became part of the top signature components of G2, G4, and G5 clusters distinctly altered in siCD81- and siCD44-transfected cells (**Figure 2A-B**), indicating one of the central pathways distinctly regulated by CD44 (RAB4A, RAB11A, RAB11B, SNX12, SNX4, and SNX6) and by CD81 (VPS29, SNX3, SNX1, SNX2, CAV2, and RAB7A) (**Supplementary Table S3, Excel Data 3**). It is well known that exosomes are generated through the endocytic pathway connected with lysosomes and exocytosis.

Interestingly, the alterations in endocytosis pathway components became more evident when phosphoproteome data were analyzed among three groups of cell lysates. Endocytosis was the top altered pathway within four out of six phosphoproteome clusters (P2, P3, P5, P6), covering both shared and distinct signaling components altered by siCD44 and siCD81. For example, both KDs promoted phosphorylation in the same residues of RAB11FP1, RAB11FP5, EPN2, and SNX12; siCD44 specifically downregulated phosphorylation in RAB8A and PLD2 but upregulated phosphorylation in GBF1, VPS26A, SNX4, RAB11FIP1, RAB11FIP5; whereas siCD81 downregulated phosphorylation in CAV2, ARFGAP1, and RAB11FIP5 (**Figure 2C-D, Supplementary Figure S3C-D, Supplementary Table S3, Excel Data 4**). These phosphoproteins regulate membrane trafficking, endosomal recycling, and caveolar formation, revealing previously unreported signaling components shared and distinguished between CD44 and CD81.

By combining 11 algorithms for machine learning-based kinase prediction, we identified the top candidates of upstream kinases potentially responsible for siCD81 and siCD44-induced alterations in phosphoproteome clusters (P1-P6), especially shared Cluster 4 with decreased phosphopeptides catalyzed by CDK1, CDK2, and EGFR, and Cluster P3 with upregulated phosphorylation by CSNK2A1 (Casein Kinase 2 α1 subunit) in (**Supplementary Figure S3E**). Furthermore, the kinase reactome analyses identified the phosphoproteome kinase networks shared between CD81 and CD44 regulation, including PAK2, one of our previously identified targets of CD44 in tumor cluster formation (36) (**Supplementary Figure S4, Excel Data 4**). These data are consistent with our previous discoveries on EGFR and PAK2 which are two representative components of the CD44 signaling and functional regulation in tumor cell clustering and metastasis (36, 61).

From an independent proteome study using the LC/MS/MS profiles of adherent MDA-MB-231 cells, the GO Processes analysis of upregulated proteins in CD44KO cells indicates regulation of proteolysis and exocytosis (**Supplementary Figure S5A, Supplementary Table S4**), whereas the GO Localization analysis of the downregulated proteins in CD44KO cells revealed the top hits of altered proteins in extracellular exosome/vesicle/organelle, previously unknown to be associated with CD44 functions (**Supplementary Figure S5B**), validating the phosphoproteome pathway regulations by CD44 and CD81 in membrane trafficking, endocytosis, lysosomes, and exocytosis in Figure 2. Thus, we hypothesized that CD44 and CD81 regulate exosome biogenesis production.

### Exosomal CD44 and CD81 promote mammosphere formation

CD81 is one of the most classical markers enriched in exosomes and/or small EVs (46, 47); however, its functions in exosomes/EVs are relatively unknown. Based on our findings that CD81 and CD44 share many signaling components regulating endocytosis and membrane trafficking, we continued to investigate whether CD44 or CD81 regulates exosome biogenesis and/or functions.

Using ultra-high resolution transmission electron microscopy, we discovered enlarged multivesicular bodies (accumulated endosomes) and increased vacuoles in both CD44KO and CD81KO cells in comparison to WT control of MDA-MB-231 cells (**Figure 3A-B**). CD81KO and CD44KO cells secreted a higher number of EVs (particles per cell) to the culture supernatants than the WT cells, as measured by the micro flow vesiclometry (MFV) as we previously established (62, 63) (**Figure 3C**). After purified from the culture supernatants of CD44KO and CD81KO cells via ultracentrifugation (**Supplementary Fig S6A**), the sizes and yield of exosome-enriched EVs (ev44KO and ev81KO) were relatively comparable to the WT control, as characterized by nanoparticle tracking analysis (NTA) and immunoblotting with exosome markers (**Figure 3D-E**, **Supplementary Figure S6B**). However, when examined by cryo-EM, ev81KO displayed impaired membrane integrity (**Figure 3F-G**), indicating an essential role for CD81 in modulating exosome biogenesis and packaging of membrane proteins.

**Figure 3.**
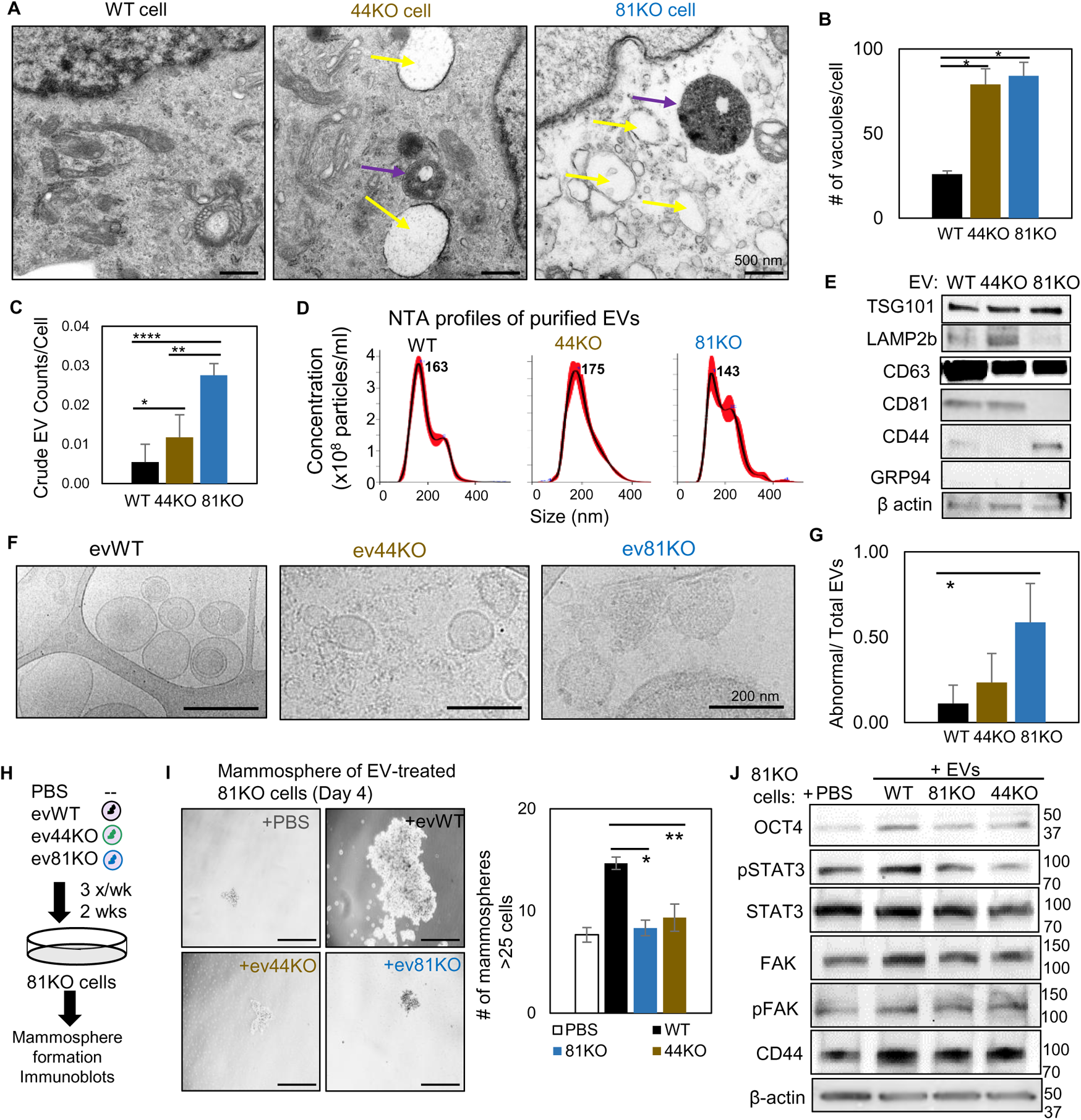
CD81 and CD44 are required for exosome-induced cancer stemness. **A-B.** Transmission electron microscopy images of WT, 44KO, and 81KO MDA-MB-231 cells (A) and number of vacuoles observed per cell (B). Yellow arrows point to vacuoles, purple arrows to multivesicular bodies. Scale bar = 500 nm. Two-tailed student T-test * P = 0.017. **C.** Counts of EVs per cell in crude culture supernatants of WT, 44KO and 81KO cells, measured by Apogee (N=3 or 5). Two-tailed student T-test **** P = 0.0001, ** P = 0.003, one-tailed student T-test * P=0.036. **D-E**. NTA-based size distributions (repeated 3 times) (D) and immunoblots (E) of ultracentrifuge-isolated EV particles from the culture media of WT, 44KO and 81KO MDA-MB-231 cells. **F-G.** Cryo-EM images (repeated twice) (F) and quantification (G) of membrane integrity in evWT, ev44KO, and ev81KO, taken at 8000x nominal magnification (a pixel size of 4.125 Angstrom). Vesicles assessed included 18 WT, 17 CD44KO, and 63 CD81KO. Scale bar = 200 nm. **H.** Schematic of 81KO MDA-MB-231 cells educated with PBS or evWT, ev44KO, and ev81KO every 2 days for 2 weeks and seeded at low density to evaluate mammosphere formation. **I.** Representative images and bar graph quantification of mammospheres 4 days after seeding of 1,000 cells per well (24-well plate). Scale bar = 100 µm. One-tailed student T-test * P = 0.02, ** P = 0.01. Repeated three times. **J.** Immunoblot analysis of 81KO MDA-MB-231 cells educated with PBS or EVs of WT, CD81KO, and CD44KO, and CD81KO cells.

A mass spectrometry analysis of the ev44WT and ev44KO identified 26 out of 416 exosomal proteins differentially expressed in ev44KO, including relatively decreased CD81 and syntenin-1 as well as upregulation of RAB-11B in association with exosome biogenesis (**Supplementary Table S5**). Proteomic pathway and network analyses identified the top altered signaling pathways related to cell adhesion, integrin-mediated matrix adhesion, cell cycle, and cytoskeleton regulation and rearrangement, and a network linking to integrins and focal adhesion (**Supplementary Figure S6C-H**), which may contribute to CD44-mediated functions in tumor metastasis (36, 61).

Next, we investigated whether, upon cellular uptake, cancer exosomes could rescue any self-renewal defects of CD81KO recipient cells, such as mammosphere formation. To do this, CD81KO MDA-MB-231 cells were educated with phosphate-buffered saline (PBS) or exosome-enriched EVs (evWT, ev44KO, and ev81KO) for 2 weeks (**Figure 3H**). Following education, the cells were evaluated for self-renewal related properties. The cells educated by evWT formed larger mammospheres with elevated protein levels of OCT4, pSTAT3, and FAK than the cells treated with PBS, ev44KO, or ev81KO (**Figure 3I-J**), demonstrating that exosomal CD44 and CD81 are required to promote self-renewal of recipient cells. Consistently, the Cd81KO 4T1 cells restored mammsphere formation after exosome education with evWT whereas no rescue effects were observed from ev81KO isolated from 4T1 cells (**Supplementary Fig S7A-B**), demonstrating Cd81-dependent regulation of self-renewal induced by mouse TNBC exosomes.

### CD81 is enriched in human CTCs and promotes CTC cluster formation

Our previous work demonstrated that CD44 mediates tumor cell aggregation and CTC cluster formation that promotes metastasis (36, 61), and is associated with reduced progression-free survival (34, 35). To determine if CD81 regulates breast cancer metastasis like CD44, we assessed the clinical relevance of CD81 expression in human breast tumors and CTCs. Public database analyses revealed that high expression of CD81 protein in breast tumors was associated with an unfavorable overall survival, relapse-free survival, and distant metastasis-free survival in patients with TNBC (**Figure 4A-C**).We also conducted a primary breast tumor TMA study and observed a correlation of CD44 and CD81 expression across tumor subtypes (TNBC, HER2, luminal A/B) as well as an upregulated expression of CD44/CD81 in TNBC in comparison to luminal A/B (**Supplementary Figure S8A-D**).

**Figure 4.**
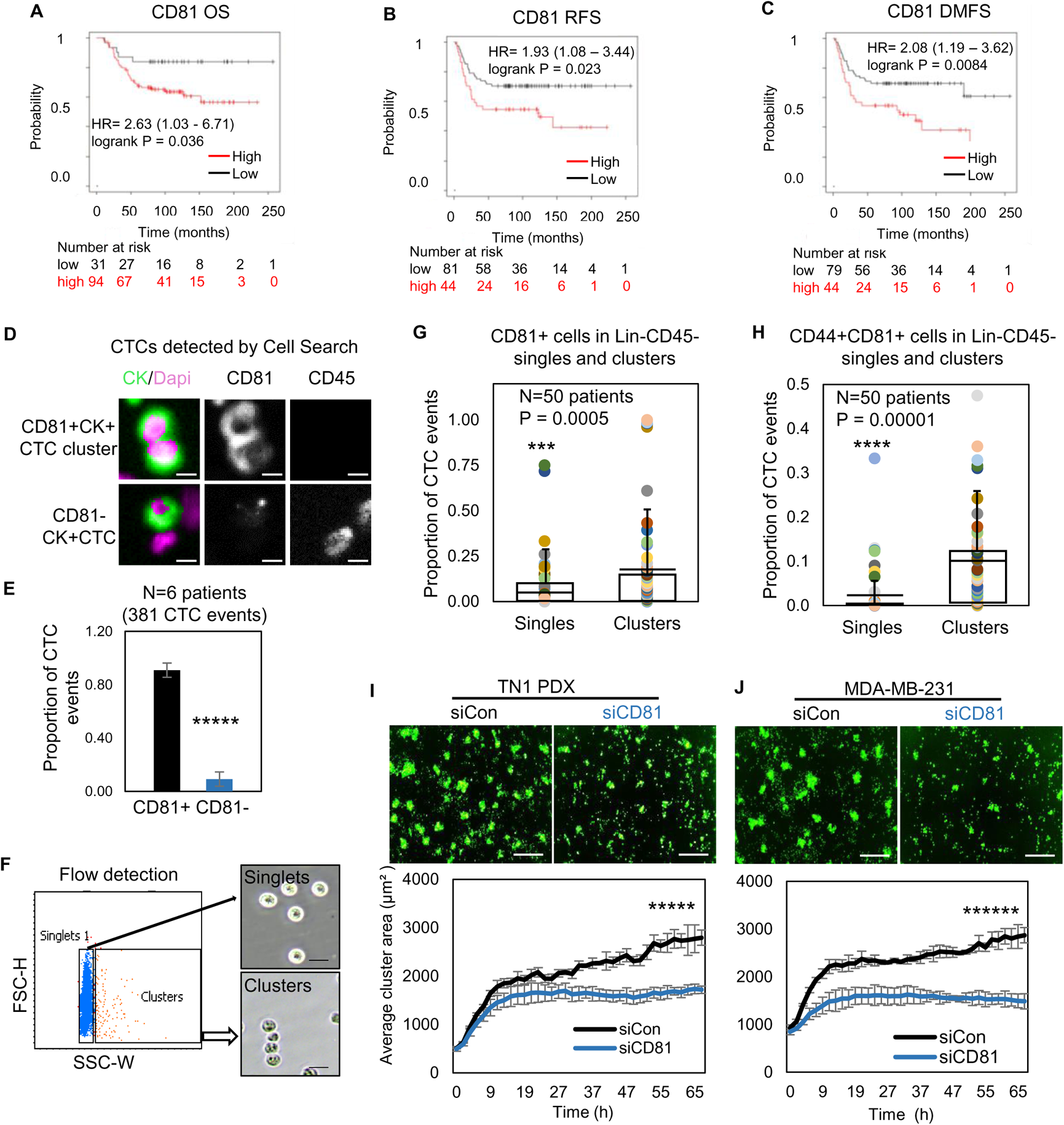
CD81 is associated with patient survival and enriched in CTCs promoting tumor cell aggregation. **A-C.** Kaplan-Mier plots of CD81 protein expression in patients with triple negative breast cancer (TNBC) correlate with an unfavorable overall survival (A), relapse-free survival (B), and distant-metastasis free survival (C). **D, E.** Representative images (D) and quantified % (E) of CD81+ and CD81- CTC events in the blood of 6 patients with metastatic breast cancer, analyzed on CellSearch. Scale bar = 5 µm. Two-tailed student T-test ***** P = 9E-11. **F-H.** Representative images of flow cytometry gated singlets and clusters (F, scale bar=25 µm) and bar graphs of proportion of CD81+ (G) and CD44+CD81+ (H) in putative Lin-CD45-CTC events (622,509) in the blood (N=50 patients). Two-tailed student T-test ***P= 0.005, ****P=0.00001. **I-J.** IncuCyte images (top panels) and quantified tumor cell aggregation curves of TN1 PDX (I), MDA-MB-231 (J) cells upon CD81 KD (repeated at least 3 times). Scale bar = 300 µm. Two-tailed student T-test ***** P = 1E-12, ****** P = 8E-19.

We then utilized three methods, immunofluorescence staining via CellSearch, flow cytometry, and RNA sequencing data analysis to further examine the CD81 expression in the CTCs isolated from patients with breast cancer. First, the FDA-approved CellSearch platform was employed to enrich EpCAM^+^ cells via anti-EpCAM beads and conduct immunofluorescence staining for validation of CD45^−^cytokerin^+^DAPI^+^ CTCs. CellSearch-based analyses of patient blood samples revealed CD81 expression in over 90% of CTC events (N=6 patients with 381 CTC events) (**Figure 4D-E**) and 100% of CTC clusters (N= two patients with 10 clusters). To expand the CD81 analysis in EpCAM^+/−^ putative CTCs, we established a flow cytometry approach to gate single cells and clusters based on size channels (forward scatter and side scatter) as validated on clustering WT and non-clustering CD44KO tumor cells (**Supplementary Figure S9A**). Consistently, flow cytometry-based analyses of putative CTCs (lineage^−^CD45^−^EpCAM^+/−^) and CTC clusters showed a significant higher expression of CD81 and CD81/CD44 double-positive expression on the clusters compared to single cells (N=50 patients, P=0.0005) (**Figure 4F-H**, **Supplementary Figure S9B**), similar to the CD44 expression enriched in CTC clusters (**Supplementary Figure S9C**) (36), suggesting a possible positive feedback loop between CD44 and CD81. Using publicly available datasets on single-cell RNA sequencing of Parsotix-filtered CTCs from the blood of patients with breast cancer (33, 35), we also found elevated CD81 expression in CTC clusters compared to single CTCs, confirming the clinical relevance of CD81 as a possible biomarker in CTC clusters and breast cancer metastasis (**Supplementary Figure S9D**).

To further determine the role of CD81 in CTC clustering, we utilized TN1 and TN2 PDX tumor cells, MDA-MB-231 cells, and 4T1 cells to analyze the phenotypic changes caused by siCD81/siCd81-mediated down-regulation or CRISPR-Cas9-mediated gene depletion. CD81 depletion or down-regulation resulted in compromised cluster formation in all 4 tested human and mouse TNBC models (**Figure 4I-J**, **Supplementary Figure S10A-C**), suggesting an important role of CD81 in promoting tumor cell aggregation. Consistently, an anti-CD81 activating agonist promoted breast tumor cell clustering in a CD81- and CD44-dependent manner as the cluster-enhancing effects of the antibody diminished in both CD81KO and CD44KO cells (**Supplementary Figure S10D**), further demonstrating the interplay and cross-dependence between CD81 and CD44 for optimal cluster formation. In addition, we performed a scratch wound assay with and without Matrigel coverage to evaluate cell invasion and migration, respectively. While CD81KO tumor cells closed the wound at a slightly slower migration speed in the absence of Matrigel compared to that of the WT control, both CD81KO cells and CD44KO cells showed much more similar and more dramatic reduction in cell invasion (**Supplementary Figure S11A-B**). We proposed to determine the role of CD81 in tumorigenesis and metastasis in the next experiment.

### CD81 promotes tumorigenesis and lung metastasis of TNBC

We first examined the importance of human CD81 and mouse Cd81 in tumorigenesis of TNBC. After dissociation, CD81^+^ and CD81^−^ TN1 PDX tumor cells were sorted on a fluorescence-activated cell sorter and then injected at dilutions of 1000- and 100-cells per injection into the mammary fat pads of NSG mice (n=4 injections/group). Up to 45 days after injection, we observed compromised tumor initiation and growth in the CD81^−^ cells, especially at the 100-cell dilution, compared to the CD81^+^ cells that grew tumors at both dilutions (**Figure 5A-C**). Similar tumorigenesis data were observed in Cd81KO 4T1 cells following 1000- and 100-cell injections (**Figure 5D-E**) (n=8 injections/group). Furthermore, CD81KO MDA-MB-231 cells had compromised tumor growth (weight) compared to WT controls when orthotopically implanted at 100 cells per mouse mammary fat pad injection (**Figure 5F-G**) (n=12 injections/group). These data demonstrate that CD81 is a newly identified promoter of breast tumor initiation.

**Figure 5.**
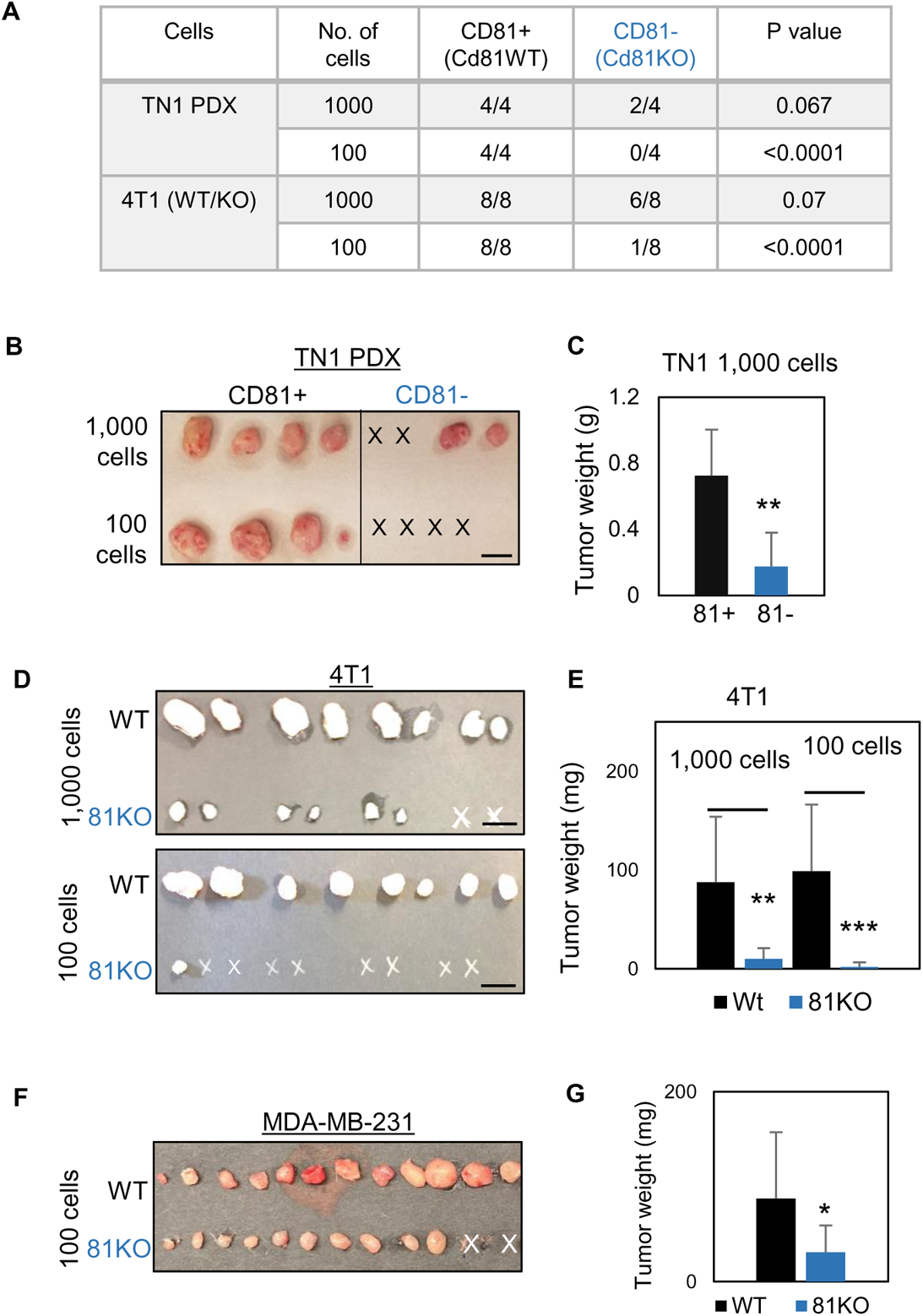
CD81 promotes tumorigenesis of TNBC cells. **A.** Table of serial dilutions of tumorigenic results with CD81+ and CD81-TN1 PDX and CD81 WT/KO 4T1 cells. One-tailed student T-test. **B-C.** Pictures of harvested tumors (B) and graphs of tumor weight comparisons (C) with CD81+ and CD81-TN1 PDX tumor implants. Scale bar=1.3 cm. n=4 injections. Two-tailed student T-test ** P=0.007 **D-E.** Pictures of harvested tumors (I), graphs of tumor weight comparisons (D) and immunoblots of 4T1 cells (Cd81 WT and KO) used for tumor implants (E). Scale bar=1 cm. n= 8 injections (2 injections/mouse). Two-tailed student T-test ** P = 0.002, *** P = 0.001. **F-G.** Pictures of harvested tumors (F) and graphs of tumor weight comparisons (G) with CD81 WT/KO MDA-MB-231 cells. Scale bar= 1 cm. n=12 injections (4 injections/mouse). Two-tailed student T-test *P= 0.016.

We continued to determine if CD81 drives spontaneous lung metastasis *in vivo*. Considering a slightly decreased tumor growth rate in mouse Cd81KO 4T1 cells, we implanted 1,000 WT cells and 6,000 Cd81KO 4T1 cells into the mammary fat pads of Balb/c mice to achieve comparable tumor burden on Day 52 when tumors and lungs were harvested (**Figure 6A-B**) (n=5 mice/group; 2 injections/mouse). While there were no significant differences in breast tumor weight between the two groups, Cd81KO tumor cells failed to metastasize to the lungs, with a significantly lower number of metastatic colonies than those of WT tumors (**Figure 6A-C**). Furthermore, the mice bearing Cd81KO tumors had 100% survival compared to 0% survival of the mice with oversized WT tumors by Day 52 (**Figure 6D**).

**Figure 6.**
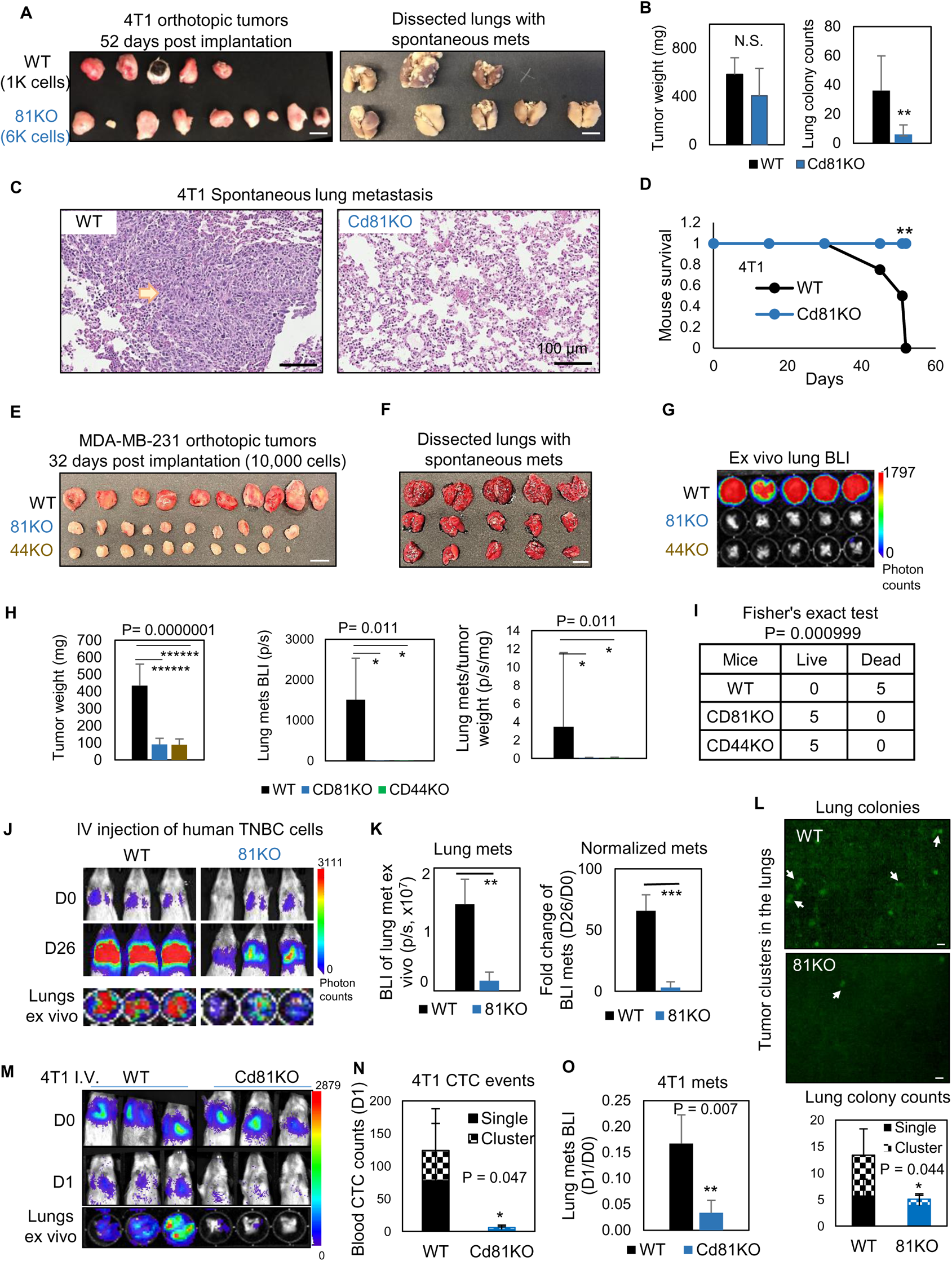
CD81 deficiency abrogated lung metastasis in breast cancer. **A.** Photos of 4T1 orthotopic tumors (left panels) grown from implantations of comparable (1000) WT and (6000) Cd81KO cells into the L4/R4 mammary fat pads (N=5 balb/c mice with 10 injections) and the fixed lungs with overt metastatic colonies. By the terminal day 52, 2 mice from the WT group died and the left 3 were sick and sacrificed. Scale bar = 1 cm. **B.** Bar graph of the tumor weights and lung colonies count between the WT tumors and KO tumors. **C.** IHC (HE) images of lung colonies from the WT group mice (a higher number of visible lung metastases at a larger size) as compared to the Cd81 KO group. Scale bar = 100 µm. N.S. = not significant, two-tailed student T-test ** P = 0.01 (n=5 mice). **D.** Distinct mouse survival between 4T1 WT and Cd81 KO tumor bearing mice with spontaneous lung metastases. Two-tailed student T-test ** P = 0.01 (n=5 mice) **E-F.** Photos of WT and 81KO MDA-MB-231 orthotopic tumors (E) grown from 10,000 cell implantations and dissected mouse lungs on day 32 (F) (N=5 NSG mice with 10 injections) Scale bar = 1 cm. **G.** BLI images and of spontaneous metastases in the lungs *ex vivo* following after orthotopic implantation of WT and 81KO MDA-MB-231cells into NSG mice. * P = 0.011 (n=5 mice). **H.** Quantification of tumor weights, lung metastases, and relative metastatic burden normalized by tumor weight. Two-tailed student T-test was used. **I.** Table of mouse mortality or survival by day 32 after orthotopic implantation. 3 mice from the WT group died and the left 2 were sick and sacrificed whereas the 81KO tumor-bearing mice would have survived. Fisher test was used ** P = 0.01 (n=5). **J.** BLI images of lung colonization following the tail vein injections of MDA-MB-231 WT and 81KO cells into NSG mice on days 0 and 26. The bottom row shows dissected lungs ex vivo. **K.** Quantified BLI signals (left panel) and normalized metastasis intensity (relative to the Day 0 signals) of MDA-MB-231 cells in the dissected lungs *ex vivo* on day 26 after tail vein injections. Two-tailed student T-test **P < 0.007. *** P = 0.001 (N=3 mice). **L.** Representative fluorescence images (top two panels) of mouse lungs and quantified metastatic colonies (singles and clustered, bottom panel) of WT and CD81KO L2G^+^ MDA-MB-231 cells (D26). Scale bar = 100 mm. Two-tailed student T-test was used. **M-O.** BLI images of mice (days 0 and 1) and dissected lungs on day 1 (M), blood CTC counts (L2G^+^ singles and clusters) measured via flow cytometry on day 1 (N), and relative lung metastasis via BLI on day 1 (O, relative to day 0) following the tail vein injections of 4T1-WT and Cd81KO cells into Babl/c mice (N=3). Two-tailed student T-test was used.

Furthermore, when human TNBC cells were implanted orthotopically at 10,000 cells to ensure tumor growth, CD81KO cells phenocopied CD44KO cells with impaired or lost capability to develop spontaneous lung metastases in mice after being normalized by tumor burden (**Figure 6E-H**) (n=5 mice/group; 2 injections/mouse), resulting in a significantly better survival of the mice bearing KO tumors than those bearing WT tumors (**Figure 6I**). Consistently, diminished lung colonization or experimental metastasis were observed in the NSG mice 26 days after receiving CD81KOcells in comparison with that of WT cells via tail vein injection, as measured via bioluminescence imaging, fluorescence microscopy, and HE-staining of lung tissues (**Figure 6J-L**, **Supplementary Fig S11C**) (n=3 mice/group). Meanwhile, siCD81 transfection-mediated transient KD of CD81 also reduced the metastatic potential of MDA-MB-231 cells following tail vein injection (**Supplementary Fig S11D-E**) (n=3 mice/group).

Finally, we measured the CTC events in the blood within one day following tail vein injection of 4T1 tumor cells and found that Cd81 KO cells (singles and clusters) were less detectable than the WT controls, in parallel with reduced seeding for lung colonization (**Figure 6M-O**) (n=3 mice/group).

In a summary, CD81 is a novel partner interacting with CD44 in breast tumor initiating cells and promotes exosome-induced self-renewal (mammosphere formation and signature markers), tumor cluster formation, and therefore enhancing tumor initiation and lung metastasis of TNBC with an unfavorable overall survival and metastasis-free survival (**Supplementary Figure 12**).

## Discussion

Our study discovers a new role of CD81 as a partner of CD44 in self-renewal related mammosphere formation, quality control of cancer exosome biogenesis, exosome-enhanced self-renewal of recipient cells, CTC cluster formation, and lung metastasis in TNBC in close association with clinical outcomes. While CD81 function has primarily been studied in immune cells (64–67), this newly identified function of CD81 in cancer metastasis is tumor cell-intrinsic and can be independent of adaptive immunity, as shown in both immunocompetent and immunocompromised mice.

Tetraspanin proteins, such as CD81, are best known for making up tetraspanin enriched microdomains (68, 69). These microdomains are crucial in regulating the motility and interactions of cancer cells with their microenvironment by organizing other transmembrane proteins, such as cell adhesion molecules, growth factors, and proteases (68, 69). To our knowledge, this is the first report linking the functions of CD81 and CD44 to endocytosis pathways and exosome-promoted mammosphere formation. Our mass spectrometry-based proteomic and phosphoproteomic profiles provide comprehensive analyses of shared and distinct signaling pathways related to CD81 and CD44 functions, including cell cycle, proliferation, and metabolism beyond exosome-related endocytosis and exocytosis. Nevertheless, other tetraspanin proteins, such as TSPAN8 and CD9, may promote cancer self-renewal in colorectal cancer (70) and glioblastoma (71), respectively. CD44 can also interact with other transmembrane proteins such as TM4SF5, resulting in elevated properties of self-renewal and circulating capacity in hepatocarcinoma cells (72).

CD44 is a multifunctional class I transmembrane glycoprotein and is widely used as a marker of breast tumor initiating cells, especially in TNBC (24). While our previous work demonstrated that CD44 homophilic binding mediates tumor cell aggregation (36, 53), in this study we propose that CD81 interacts with CD44 to promote intracellular CD44-CD81 heterodimer formation and possibly intercellular tetramer formation between two neighboring cells. CD44 and CD81 interactions on the cell membrane might provide feedback for protein networks and modifications. Notably, a transmembrane ubiquitin ligase family member MARCH8 has been associated with lysosome degradation of both CD44 and CD81 in fibroblast cells (73) and/or TNBC cells (74), suggesting CD44 and CD81 might follow similar fates of protein degradation or recycling in both cancer cells and other cells. Nevertheless, both CD44 and CD81 are required for optimal self-renewal and metastasis, emphasizing the indispensable functions of CD44 and CD81 in cell adhesion and intercellular interactions in metastasis. Our phosphoproteome analyses reveal many shared and unshared components between CD44 and CD81 signaling pathways that regulate endocytosis, endosomes, lysosomes, and exocytosis.

Several studies suggest tumor recurrence often occurs due to an increased number of CTCs, some of which display tumor-regenerative plasticity and reprogramming phenotypes (75–77), or transform into tumor-initiating cells (78–81). Our studies demonstrated that cancer exosomes can be part of the transforming factors upon uptake by the CTCs. Here we show that the mammosphere-promoting functions of exosomal CD44 and CD81 illustrate the crosstalk between tumor-initiating cells and surrounding cancer cells that potentially contributes to their self-renewal and/or plasticity. While CD44 remains understudied in exosome biogenesis, a CD44 variant has been reported to be involved in interluminal vesicle loading (82) which may be linked to maintaining tumor-initiating cells and tumor progression. Furthermore, a subset of CTCs and tumor-initiating cells may exhibit dynamic changes in epithelial–mesenchymal transition (EMT) phenotype (83), which can up-regulate CD81 expression in mesenchymal breast cancer (84). While the role of CD81 displayed on CTCs is largely understudied, previous studies have used CD81^+^/CD56^+^/CD45^−^ markers to detect neuroblastoma cells in the peripheral blood of patients (85). Future studies will be needed to investigate the effects of cancer exosomes on tumor stromal cells and immune cells in various microenvironment niches, not only in TNBC, but also in other cancers as well.

## Methods

### Human specimen analyses

All human blood and tumor specimen analyses complied with NIH guidelines for human subject studies and were approved by the Institutional Review Boards at Northwestern University. The investigators obtained written informed consent from all subjects whose blood specimens were analyzed.

### Animal studies

All mice used in this study were kept in specific pathogen-free facilities in the Animal Resources Center at Northwestern University. All animal procedures complied with the NIH Guidelines for the Care and Use of Laboratory Animals and were approved by the respective Institutional Animal Care and Use Committees. Animals were randomized by age and weight. Mice were excluded from experiments for sickness or conditions unrelated to tumors. Sample sizes were determined based on the results of preliminary experiments, and no statistical method was used to predetermine sample size. All PDX tumors were established, and orthotopic tumor implantation was performed as described previously (16, 86). For tumorigenic assays, cells and Matrigel (Corning 354234) were implanted into the mammary fat pad of mice at low density (1000-100 cells). Mice were monitored using the Lago *in vivo* imaging system. For artificial metastasis experiments, (100,000) tumor cells were injected into the mice tail vein and imaged using the Lago *in vivo* imaging system. For spontaneous metastasis experiments using MDA-MB-231 cells, 10,000 cells were implanted orthotopically in NSG mice. For spontaneous metastasis experiments using 4T1 cells, 1,000 WT cells and 6,000 Cd81KO cells were implanted orthotopically in Balb/c mice.

### Blood collection and CTC analysis

This study was approved by the Institutional Review Board (IRB) of Northwestern University. All participants signed informed consent forms. About 8-10 ml of whole blood was collected from breast cancer patients into a 10 ml CellSave Preservative tube containing a cellular fixative (Janssen Diagnostics, LLC, Raritan, NJ). Blood specimens were maintained at room temperature and processed within 96 h of being drawn. CTC analysis was performed using CellSearch® CTC kits on the FDA-approved CellSearch System (Item 7900001, Menarini Silicon Biosystems, Inc). The kit uses ferrofluid nanoparticles with antibodies that target epithelial cell adhesion molecules (EpCAM) to magnetically separate CTCs from the bulk of other cells in the blood. CTCs were identified by positive staining for both cytokeratins (CK) and DAPI and negative staining for CD45 (CK+/DAPI+/CD45-). CTC clusters were defined as an aggregation of two or more CTCs containing distinct nuclei and intact cytoplasmic membranes. To determine the expression of CD81 on CTCs, the PE-conjugated anti-CD81 antibody (BD) was also added. CTCs were also analyzed using flowcytometry by gating single cells and cell clusters for CD45 negative and DAPI negative. The proportion of CTC events was calculated by dividing the population of interest by the sum of total CTC events. For example, to calculate the percentage of CD44+CD81+ CTCs in a patient, CD44+CD81+ events were divided by the sum of CD44+CD81+, CD44+CD81-, CD44- CD81+, and CD44-CD81-events.

### Cell culture, CRISPR gene knockout, and transfections

MDA-MB-231 and HEK293ft cells were purchased from ATCC, and periodically verified to be *Mycoplasma*-negative using MycoAlert Mycoplasma Detection Kit (Lonza cat #LT07-218). Cell morphology, growth characteristics, and microarray gene-expression analyses were compared with published information to ensure their authenticity. Early passages of cells (<20 passages) were maintained in Dulbecco’s Modified Eagle Medium with 10% fetal bovine serum (FBS) + 1% penicillin–streptomycin (P/S). Pooled populations of wildtype (WT) control cells, CD81KO MDA-MB-231 cells, and Cd81KO 4T1 cells were made using multiple lentiviral gRNAs with a BFP reporter for each gene (Sigma Cat# HS5000016583 and HS5000016584 for human *CD81*, MM5000007469 and MM5000007470 for mouse *Cd81*, and gRNA control vector) and CRISPR-Cas9 with a GFP reporter (Sigma Cat# CMV-CAS9-2A-GFP), and flow sorted based on expression of GFP and BFP reporters and absence of CD81/Cd81. CD44KO cells were generated based on the validated lentivral gRNAs and protocol described previously (36) and then combined with CD81KO for generation of dKO cells (pooled populations).

For exosome experiments, FBS was exosome-depleted by ultracentrifugation at 100,000 × g for 16 h at 4 °C. Primary tumor cells were cultured in HuMEC-ready medium (Life Technologies) plus 5% FBS and 0.5% P/S in collagen type I (BD Biosciences) coated plates. Pooled siRNAs (SMART pool with 3 to 4 siRNAs; Dharmacon CD81 Cat# L-017257-00, Dharmacon negative control non-targeting pool Cat# D-001810-10-50) were transfected using Dharmafect (Dharmacon) at 100 nmol/L. For overexpression experiments in HEK293ft cells, pCMV6-FLAG-CD44 (OriGene) and pCMV3-HA-CD81 (Sino Biological) plasmids were transfected into cells using Fugene HD (Promega). After 48 h, cells were collected for immunoprecipitation and immunoblotting.

### Mammosphere formation, cell clustering, and migration assays

For mammosphere assays, cells were seeded into 6-well or 12-well plates at a concentration of 2,000 or 1,000 cells per well in replicates of 3 or 4 using Prime-XV Tumorsphere Serum Free Media (Irvine Scientific). Cells were monitored for up to 17 days, when spheres were imaged and counted to assess mammosphere formation capacity. For exosome education experiments, cells were seeded into 12-well plates at a concentration of 50,000 cells and treated with 10-15 ug exosomes every other day for 1-2 weeks. The cells were split as necessary and mammosphere formation assay followed (2,000 cells seeded per well). For clustering assays, cells were seeded into polyhema-coated 96-well plates at a concentration of 25,000 cells per well for cell lines and 100,000 cells per well for primary cells. Cells were monitored up to 72 h and analyzed by Incucyte Imaging System software. For migration assays, cells were seeded into 96-well Image-Lock plates at 30,000 cells per well. After 16 h, a scratch wound was introduced into each plate using the Incucyte Wound Maker and then monitored by Incucyte for up to 48 h to visualize wound closure.

### Immunofluorescence

Cells were seeded onto 4-chamber slide wells at a concentration of 20,000 cells per well. Clustered cells were attached using a cytospin. Cells were fixed with 4% paraformaldehyde and permeated with 1% Triton X-100 in PBS for 15 min at room temperature. Cells were then washed 3 times for 5 min each with 0.05% PBS and blocked with 10% bovine serum albumin (BSA) in PBS for 60 min. Cells were then washed again and primary antibody was added overnight at 4 °C (CD81 Millipore HPA007234 4 µg/ml and CD44 MA513890 2 µg/ml). After washing, secondary antibody was added for 60 min at room temperature (Texas Red T862 Thermo-Fisher and Alexa-488 A11008 Thermo-Fisher). Finally, the cells were washed, and a cover slide was placed with mounting media. The slides were imaged using Nikon A1R (A) Spectral.

### Structural modeling

The structure of CD44 (the shortest standard isoform X4) was predicted by the iTasser webserver (55), with abundant homologous structures available to the N-terminal extracellular domains. And the structure of CD81 in a “closed” conformation was obtained from the Protein Data Bank (PDB ID: 5TCX), with a missing extracellular loop (residues 38-54) inserted (54). The two structures were first rigidly docked while being biased toward extracellular regions, using the ClusPro webserver (56). The resulting structural models of CD44-CD81 complexes were then flexibly refined, using the software Bayesian Active Learning (BAL) (52), where binding hotspots (in probabilities from 0 to 1) and binding affinity (in kcal/mol) are predicted and weighted-averaged over all structural models. Two short extracellular helices (resi. 160-170 and 181-187) were predicted to be enriched in binding hotspots and their borders include C156 and C190 forming a disulfide bond. Therefore, a form of CD81 with S159-K187 deletion (CD81d) was suggested to impair its interaction with CD44 while maintaining its stability. The structure of CD81d was again predicted with iTasser and the docking and analyses of CD44/CD81d followed the protocol described above.

### Flow cytometry and cell sorting

To detect cell surface proteins, cells were first blocked with mouse IgG 1 (Cat# l5381, Sigma, St. Louis, MO, USA) for 10 min on ice. Cells were then incubated with antibody (BD Biosciences, San Jose, CA, USA) for 20 min on ice, washed, and analyzed using a BD LSR-2 flow cytometer or BD Aria cell sorter (BD Biosciences).

### RNA sequencing

MDA-MB-231 cells were seeded and transfected with non-targeting siRNA (siCon) and siCD81 (pooled, Dharmacon) using Dharmafect. After 48 h, the cells were harvested and submitted for RNA sequencing. Total RNA of MDA-MB-231 cells was isolated using Trizol, phase separated by chloroform, and extracted by alcohol. Samples were sent to Northwestern University’s Center for Genetic Medicine Sequencing core facility for deep sequencing analysis. RNA sequencing was performed on a HiSeq 4000, and a library was made using aTruSeq Total RNA-Seq Library Prepkit. Data were processed and quantified using STAR (87), DESeq2 (88), and HTSeq (89). Analysis of differentially expressed genes was set to a cutoff of false discovery rate < 0.05 and log2 (fold change) > 0.48 or < −0.48. Finally, the pathway analysis of significantly differentially expressed genes was obtained by using Metascape (http://metascape.org) (57). The raw data files of RNA seq data generated with control and siCD81-transfected MDA-MB-231 cells have been deposited to GEO database with accession number GSE174087.

### Mass spectrometry of tumor cells and exosomes

Exosomes were isolated from MDA-MB-231 WT control and CD44KO cell cultures via standard ultracentrifugation as described (63). The cells and exosomes were lysed with 2% SDS and protease inhibitor cocktail. Proteins were extracted using pulse sonication, and cleaned up by filter-aided sample preparation (FASP) to remove detergents. After LysC/Trypsin digestion, 500 ng proteins were analyzed via 4-h LC/MS/MS method at Case Western Reserve University Proteomics Core facility and the data processed using MetaCore. The fold change was calculated based on total unique spectrum counts.

### Global and phosphoproteome analyses of tumor cells transfected with siCD81 and siCD44

For protein extraction, cell pellets were resuspended in cell lysis buffer (100 mM NH_4_HCO_3_, pH 8.0, 8 M urea, 1% protease and phosphatase inhibitor, pH 8.0) and protein concentrations were measured with a Pierce BCA protein assay (Thermo Fisher Scientific). Proteins were reduced with 5 mM dithiothreitol for 1 h at 37°C and alkylated with 10 mM iodoacetamide for 45 min at 25°C in the dark. Protein was digested with Lys-C (Wako) (1:50 enzyme-to-substrate ratio) for 3 h at 25°C and with sequencing-grade modified trypsin (Promega, V5117) at 25°C for 14 h. After digestion, each sample was desalted by C18 SPE extraction and concentrated for BCA assay to evaluate the peptide yield.

The tryptic peptides from bulk samples were dissolved with 50 mM HEPES (pH 8.5) and then mixed with a TMTpro reagent in 100% ACN. A ratio of TMTpro to peptide amount of 4:1 was used. After incubation for 1 h at room temperature, the reaction was terminated by adding 5% hydroxylamine for 15 min. The TMTpro-labeled peptides were then acidified with 0.5% FA. Peptides labeled by different TMTpro reagents were then mixed, dried using Speed-Vac, reconstituted with 3% acetonitrile, 0.1% formic acid and desalted on C18 SepPak SPE columns.

Peptide fractionation by bRPLC and phosphopeptides enrichment by IMAC were performed as previously reported (90). Lyophilized global and phosphorylated peptides were reconstituted in 12 μL of 0.1% FA with 2% ACN and 5 μL of the resulting sample was analyzed by LC-MS/MS using an Orbitrap Fusion Lumos Tribrid Mass Spectrometer (Thermo Scientific) connected to a nanoACQUITY UPLC system (Waters Corp., Milford, MA) (buffer A: 0.1% FA with 3% ACN and buffer B: 0.1% FA in 90% ACN) as previously described (91). Peptides were separated by a gradient mixture with an analytical column (75 μm i.d. × 20 cm) packed using 1.9-μm ReproSil C18 and with a column heater set at 50 °C. Peptides were separated by a gradient mixture : 2-6% buffer B in 1 min, 6-30% buffer B in 84 min, 30-60% buffer B in 9 min, 60-90% buffer B in 1 min, and finally 90% buffer B for 5 min at 200 nL/min. Data were acquired in a data dependent mode with a full MS scan (m/z 350-1800) at a resolution of 60K with AGC setting set to 4×10^5^ and maximum ion injection period set to 50 ms. The isolation window for MS/MS was set at 0.7 m/z and optimal HCD fragmentation was performed at a normalized collision energy of 30% with AGC set as 1×10^5^ and a maximum ion injection time of 105 ms. The MS/MS spectra were acquired at a resolution of 50K. The dynamic exclusion time was set at 45 s. Raw data sets have been deposited in the Japan ProteOmeSTandard Repository (https://repository.jpostdb.org/) (92). The accession numbers are PXD029529 for ProteomeXchange (93) and JPST001321 for jPOST. The access link is https://repository.jpostdb.org/preview/1370203119618182ba1c0f2 with (access key 7811 for reviewer only until accepted).

The raw MS/MS data were processed with MSFragger via Fragpipe (94, 95) with TMT16 quantitation workflow. The MS/MS spectra were searched against a human UniProt database (fasta file dated July-31, 2021 with 40,840 sequences which contain 20,420 decoys). The intensities of all sixteen TMT reporter ions were extracted from Fragpipe outputs and analyzed by Perseus (96) for statistical analyses. The abundances of TMTpro were firstly log2 transformed. The TMT intensities were normalized based on the column-centering by median values for statistical pairwise comparison. For pathway analysis, the significantly expressed protein or phosphophoproteins after Anonva t-test analysis (FDR<0.01) were analyzed by DAVID (97). The protein-protein interaction analysis was performed with STRING (98) and Cytoscape (99).

MaxQuant was used to process the raw MS/MS data with 20,198 sequences recognized against a human UniProt database (fasta file dated April 12, 2017), default setting for mass tolerance for precursor and fragment ions and “Reporter ion MS2” for isobaric label measurements. A peptide search was performed with Trypsin/P and allowed a maximum of two missed cleavages. Carbamidomethyl (C) was set as a fixed modification; acetylation (protein N-term), oxidation (M) and Phospho (STY) were set as variable modifications for phosphoproteome analysis. The false discovery rate (FDR) was set to 1% at the level of proteins, peptides, and modifications. The Phospho (STY) Sites.txt file was used for further quantitation. The intensities of all ten TMT reporter ions were extracted from MaxQuant outputs and analyzed by Perseus (96) for statistical analyses.

### Isolation and purification of exosomes from cells

Exosomes were isolated from the cell culture supernatant as described previously (63). Briefly, the cells were cultured as monolayers for 48 h in complete medium under an atmosphere of 5% CO2 at 37 °C. When cells reached a confluency of approximately 80%, exosomes were isolated by differential centrifugation. First, the culture supernatant was centrifuged at 2,000 × g for 10 min followed by 30 min centrifugation at 10,000 × g to remove dead cells and cell debris. The supernatant was ultracentrifuged for 70 min at 100,000 × g using an SW28 rotor to pellet the exosomes. Exosomes were washed by resuspension in 30 ml of sterile PBS (Hyclone, Utah, USA), and pelleted by ultracentrifugation for 70 min at 100,000 × g. The final exosome pellet was resuspended in 100 μl PBS and stored at −80 °C.

### Transmission electron microscopy

Cells were harvested from the medium and washed with Na/K Phosphate buffer (0.1 M, pH 7.2). Primary fixation was done with 3% glutaraldehyde for two hours at 4 ◦C. The cells were then post-fixed in 1% osmium tetroxide for 1 h at 4 ◦C. The cells were resuspended in molten agar (2%). Small blocks of solidified agar (1sq.mm) were cut and passed through series of 30%, 50%, and 90% ethanol v/v for 15 minutes each. The cells were further dehydrated with 100% ethanol (30 min x 3). The dehydrated agar blocks were suspended in propylene oxide for 20 minutes at room temperature and then treated with 1:1 mixture of propylene oxide and Epon-812 for 1 h at room temperature and Epon-812 for 4 h at room temperature. Blocks were embedded with Epon-812 for 48 h at 60 ◦C. Ultra-thin sections were cut with a Leica UC6 ultramicrotome and examined with a FEI Tecnai Spirit transmission electron microscope (FEI, Hilsboro City, OR, USA).

### Cryo-electron microscopy

For cryoEM visualization, samples were prepared from freshly isolated exosomes at 0.25 µg/µl concentration. For cryo-freezing, 3.5μl of exosome solutions were applied to fresh glow-discharged (10 s, 15 mA; Pelco EasiGlow) lacey carbon TEM grids (Electron Microscopy Services) and vitrified using a FEI Vitrobot Mark IV (FEI, Hillsboro, OR). The sample was applied to the grid and kept at 85% humidity and 10 oC. After a 10 second incubation period the grid was blotted with Whatman 595 filter paper for 3.5 seconds using a blot force of 5 and plunge frozen into liquid ethane. Samples were imaged using a JEOL 3200FS electron microscope equipped with an omega energy filter operated at 300 kV with a K3 direct electron detector (Ametek) using the minimal dose system. The total dose for each movie was ~10 e-/A2 at a nominal magnification between 8,000 (pixel size 4.1 Å).

### NanoSight particle tracking and rapid microflow cytometer analysis of exosomes or extracellular vesicles (EVs)

The size and particle count of exosomes were measured using NanoSight NS3000, a nanoparticle tracking analyzer (NanoSight Ltd, Malvern, United Kingdom). Exosomes (5 µg) were diluted in 1 ml PBS and then processed. Similarly, direct supernatants after removal of cell debris or purified exosomes/EVs diluted in 300 µL of PBS were loaded to the Apogee micro flow vesiclometer for EV count analysis and normalized based on the cell numbers to compare the EV secretion efficiency or yield.

### Immunoblot analysis

Cells and exosomes were lysed using RIPA lysis buffer. Protein-containing lysates of exosomes (5 μg) were run on a 4–20% Mini-PROTEIN TGX gel (Bio-Rad, Hercules, California, USA) and transferred to a nitrocellulose membrane. The blots were incubated separately either with mouse monoclonal anti-human CD44 (156-3C11) antibody (Cat# MA513890, ThermoFisher Scientific, Waltham, MA, USA), mouse monoclonal anti-human CD63 antibody (Cat# ab8219, Abcam, Cambridge, MA, USA) at a dilution of 1:1000, Rabbit polyclonal anti-human CD81 (Cat# GTX101766, Genetex, Irvine, CA, USA) at a dilution of 1:1000, rabbit polyclonal anti-human Grp94 (Cat# 2104P, Cell Signaling Technology, Danvers, MA, USA) at a dilution of 1:1000, or mouse monoclonal anti-human β-actin (Cat# ab8224, Abcam, Cambridge, MA, USA) at a dilution of 1:1000 (in Tris-buffered saline (TBS) containing 2% BSA) at room temperature for 1 h, followed by washing with TBS buffer. The blots were incubated with secondary antibody (horseradish peroxidase-conjugated goat anti-mouse (Cat# W402B) or goat anti-rabbit IgG (W401B) from Promega, Madison, WI, USA) at a dilution of 1:10,000 (in 2% BSA containing TBS) for 1 h at room temperature. The blots were treated with an enhanced chemiluminescence kit according to the user manual and developed using a ChemiDoc MP Imaging System (BioRad).

### Co-immunoprecipitation

For endogenous co-immunoprecipitation, cells were lysed and preincubated with Protein A/B PLUS agarose beads (Cat# sc-2003, Santa Cruz Biotech, Dallas, TX, USA) for 2 h. The supernatant was removed, and the protein concentration was measured. Then 100 ug of cell lysate was incubated with CD44 anti-body or bead control overnight and then Protein A/B PLUS agarose beads overnight. The beads were washed 5 times and denatured with 4x Laemmli sample buffer (Cat: 161-0747, BioRad, Hercules, CA, USA) at 100 C for 5 min. For exogenous co-immunoprecipitation, cells were lysed, and protein concentration was measured. Then 200 ug of cell lysate was incubated with Anti-FLAG® M2 Magnetic Beads, Sigma-Aldrich (Cat# M8823, Sigma-Aldrich, St. Louis, MO, USA) overnight. The next day the beads were washed, and the beads were eluted using glycine. The elution was then combined with 4x Laemmli sample buffer and denatured at 100 C for 5 min.

### Immunohistochemistry

Mouse xenograft lung tissues or patient primary tumors were paraffin-embedded and sectioned by routine techniques. Heat-induced antigen retrieval was achieved using Decloaker solution for 15-20 min (Biocare Medical, RD913L). Tissue sections were blocked with TBS/10% NGS, then incubated with CD81 (Cat # HPA007234 Millipore Sigma, St. Louis, MO, USA) or CD44 (Cat# MA513890, ThermoFisher Scientific, Waltham, MA, USA) primary antibody overnight, followed by Dako envision plus kit and DAB staining. All samples were counterstained with hematoxylin.

### RNA extraction and real-time PCR

Total RNA was extracted using Trizol (Invitrogen), and RNA was paecipitated with isopropanol and glycogen (Invitrogen). After reverse transcription reactions, real-time PCR for genes was performed using individual gene Taqman primers (Applied Biosystems) with an ABI 7500 real-time PCR system. GAPDH was used as a control.

### Kaplan-Meier plots

Kaplan-Meier overall survival, relapse free survival, and distant metastasis free survival plots for protein expression of CD81 were made using kmplot.com. The dataset Liu_2014 (n=126) was used. All patients had triple negative breast cancer.

### Breast tumor tissue microarray (TMA)

A total of 89 formalin-fixed paraffin-embedded breast tumor tissues were included on the tumor TMA with selected tumor regions guided by hematoxylin-eosin staining images. The demographic and clinical characteristics of these tumors are included in Supplementary Figure S8. To make the TMA that allows microscopic comparison of the staining characteristics of different blocks and prevents exhaustion of pathological material, a core of paraffin was removed from a “recipient” paraffin block (one embedded without tissue) and the remaining empty space is filled with a core of paraffin embedded tissue from a “donor” block. A donor block H&E that is representative of the tissue remaining in the block was used to select the sample core with a color marker corresponding to tumor, benign, etc. Matched blocks were pulled out and a recipient TMA block was made and trimmed well with the face of the block even with a size of 1.5 mm core by using the semi-automatic Veridiam Tissue Microarryer VTA-100. The created TMA block was sectioned for staining. In this TMA, 19 cases from NU 16B06, 9 ER negative cases, 30 triple negative cases, 27 ER positive cases and 4 normal breast cases were selected and constructed on the recipient block.

### Statistical analysis

For all assays and analyses *in vitro*, unless otherwise specified, a two-tailed Student’s *t* test performed using Microsoft Excel was used to evaluate the *P* values, and *P* < 0.05 was considered statistically significant.

## Acknowledgements

We are grateful for the tremendous support by Northwestern University Core facilities, including but not limited to the CTC Core, the Center for Comparative Medicine, Flow Cytometry Core, Small Animal Imaging, Microscopy Imaging, NUSeq, Bioinformatics, Mouse Histology & Phenotyping Laboratory, and Pathology Core. We also thank Case Western Reserve University Mass Spectrometry Core and Cancer Center Small Animal Facilities for their support. This project has been partially supported by the Department of Defense grant W81XWH-16-1-0021 and W81XWH-20-1-0679 (H. Liu), the NIH/NCI grants R01CA245699 (H. Liu and E.K. Ramos); NIH/NIGMS R35GM124952 (Y. Shen); National Science Foundation CCF-1943008 (Y. Shen); the Lynn Sage Cancer Research Foundation (X. Liu, M. Cristofanilli, and H. Liu), Susan G. Komen Foundation CCR18548501 (X. Liu); American Cancer Society ACS127951-RSG-15-025-01-CSM (H. Liu); Northwestern University start-up grant (H. Liu), and NIH Fellowships T32 CA009560 (E.K. Ramos), T32 CA080621-15 and the Julius Kahn Fellowship (R. Taftaf), and T32GM008061 (E.J. Schuster).

---

**Suppl. Fig. S1.**
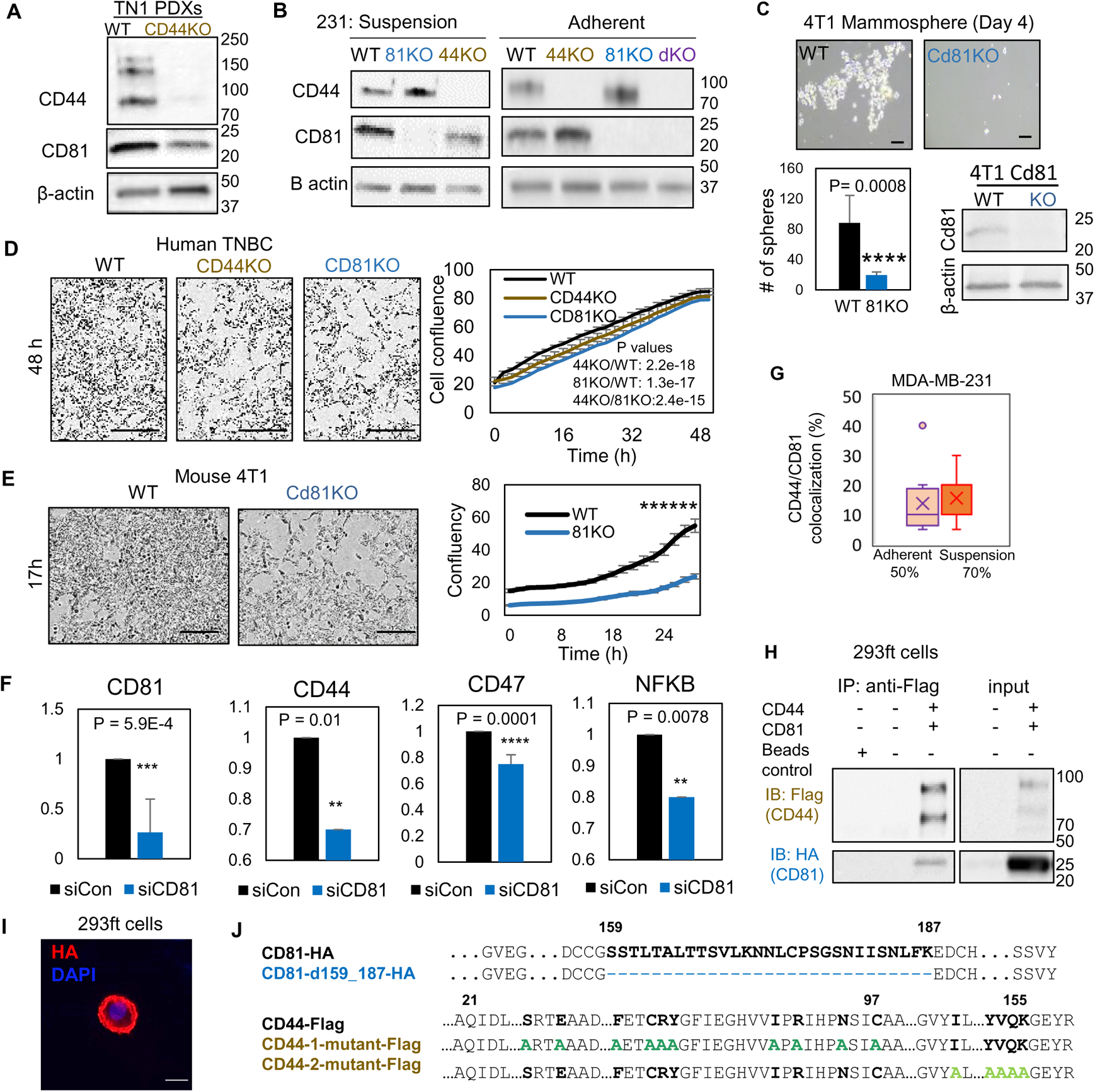
Characterization of CD44KO and CD81KO breast cancer cell lines (related to Fig. 1). **A, B.** Immunoblots of CD44 and CD81 in dissociated PDX (WT and CD44KO) tumor cells in suspension and MDA-MB-231 cells, in suspension and adherent (WT, CD44KO, CD81KO, and dKO). **C.** Mammosphere formation and immunoblot of mouse 4T1 cells (WT and Cd81 KO via CRISPR/Cas9). Cells were seeded 2000 cells/ well in 6 well plate in 4 replicate.Two-tailed student T-test was used. Repeated twice. **D-E.** IncuCyte images and curve analyses of cell confluence of human MDA-MB-231 WT, CD44KO, and CD81KO cells (C) and mouse 4T1 WT and Cd81KO cells (D) over time Scale bar is 300 µm. Two-tailed student T-test ****** P =1.97E-08. **F.** qRT-PCR expression of CD81, CD44, CD47, and NFKB in siCon and siCD81 MDA-MB-231 cells. Two-tailed student T-test *** P = 5.9E-4, * P = 0.012, **** P = 0.0001, ** P = 0.0078. Repeated 3 times. **G.** % of CD44 and CD81 colocalization shown in 50% of adherent (14 out of 26 cells) and 70% of suspension (22 out of 31 cells) MDA-MB-231 cells (WT). **H.** Immunoblots of exogenously expressed CD44-Flag and CD81-HA immunoprecipitated by anti-Flag (CD44) from the lysates of transfected HEK293ft cells. Repeated 3 times. **I.** Immunofluorescence of HA tagged CD81d in HEK293ft cells showing its membrane localization. **J.** Alignment of CD44 and CD81 amino acids showing deletion in CD81 and point mutations in CD44 domains I & II.

**Suppl. Figure S2.**
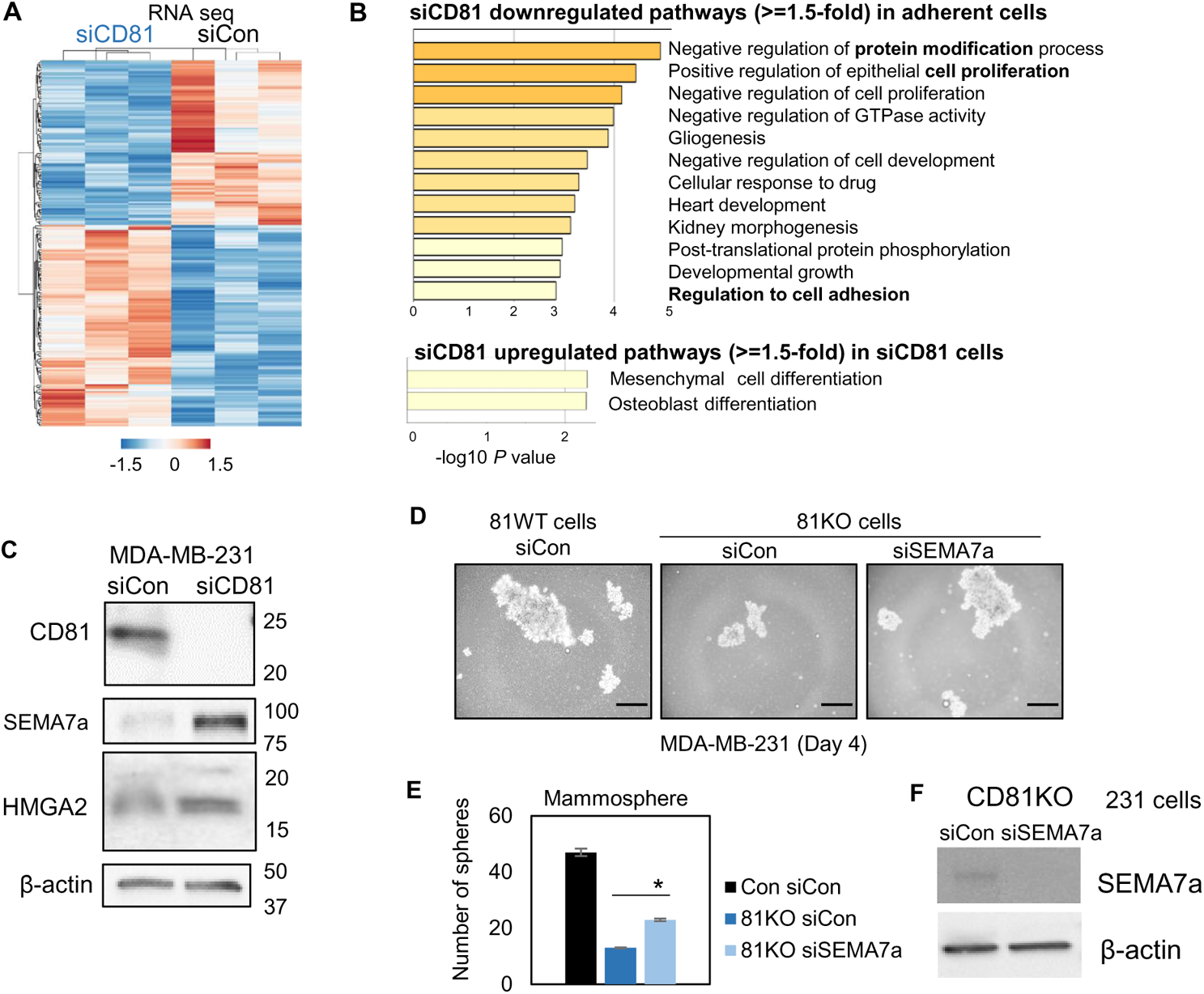
CD81 target pathways and genes (related to Fig. 2) **A, B.** RNA-sequencing based transcriptome analysis show siCD81-altered pathways (A),and heatmap (B), including downregulated and upregulated genes with >=1.5-fold changes) in MDA-MB-231 cells compared to the siRNA control (N=3 replicates). **C.** Immunoblots of top targets SEMA7a and HMGA2 altered by siCD81 in adherent cells. **D-F.** Representative images of formed mammospheres (D), quantification (E) of mammospheres, and immunoblots of SEMA7α after siSEMA7a gene KD in CD81KO cells (F), showing downregulation of Sema7a rescues mammosphere formation of CD81KO MDA-MB-231 cells, Sema7a was knocked down via siRNA transfection and seeded at 2000 cells/well in 12-well plate. After 4 days, Sema7a KD in CD81KO cells increased mammosphere number and size compared to the control. Scale bar = 50 µm. * P = 0.015.

**Suppl. Figure S3.**
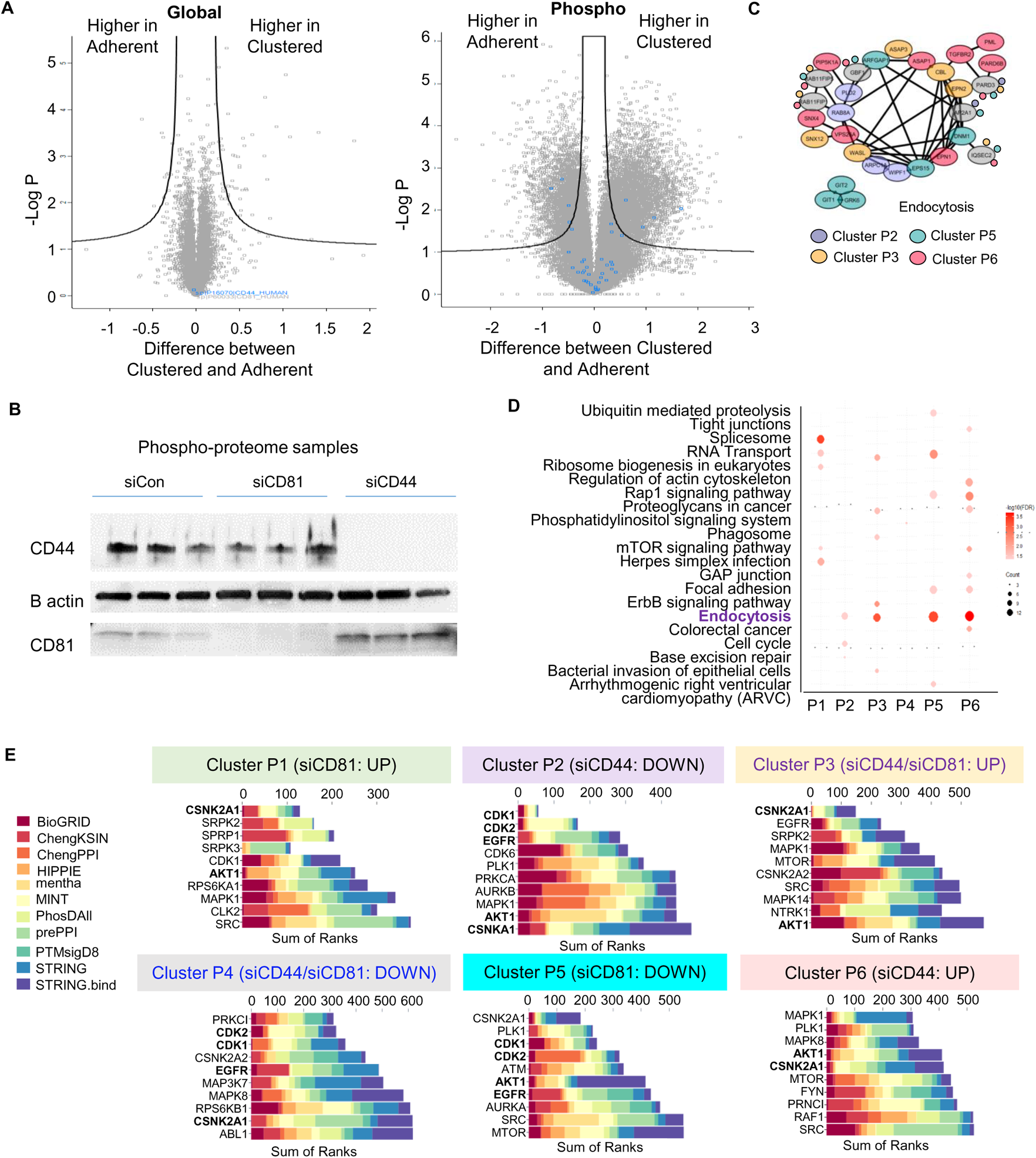
Global and phosphoproteomic analyses between adherent and tumor cells in suspension and among siControl, siCD81, and siCD44 tumor cells (related to Fig. 2) **A.** The Volcano plots of different abundance in global proteome and phosphoproteome between adherent and clustered tumor cells. **B.** Immunoblots of CD44 and CD81 in MDA-MB-231 cells after transient knockdowns after siCD81and siCD44 transfections. **C.** The protein-protein interactions network of altered phosphoproteome in the Endocytosis pathway across four phosphoproteome clusters P2, P3, P5, P6 **D. E.** The enrichment of KEGG pathway (D) and Kinase Enrichment Analysis (KEA) (E) from expressed phosphopeptides in the different clusters P1-P6 comparing siControl with siCD81 and siCD44 groups.

**Suppl. Figure S4.**
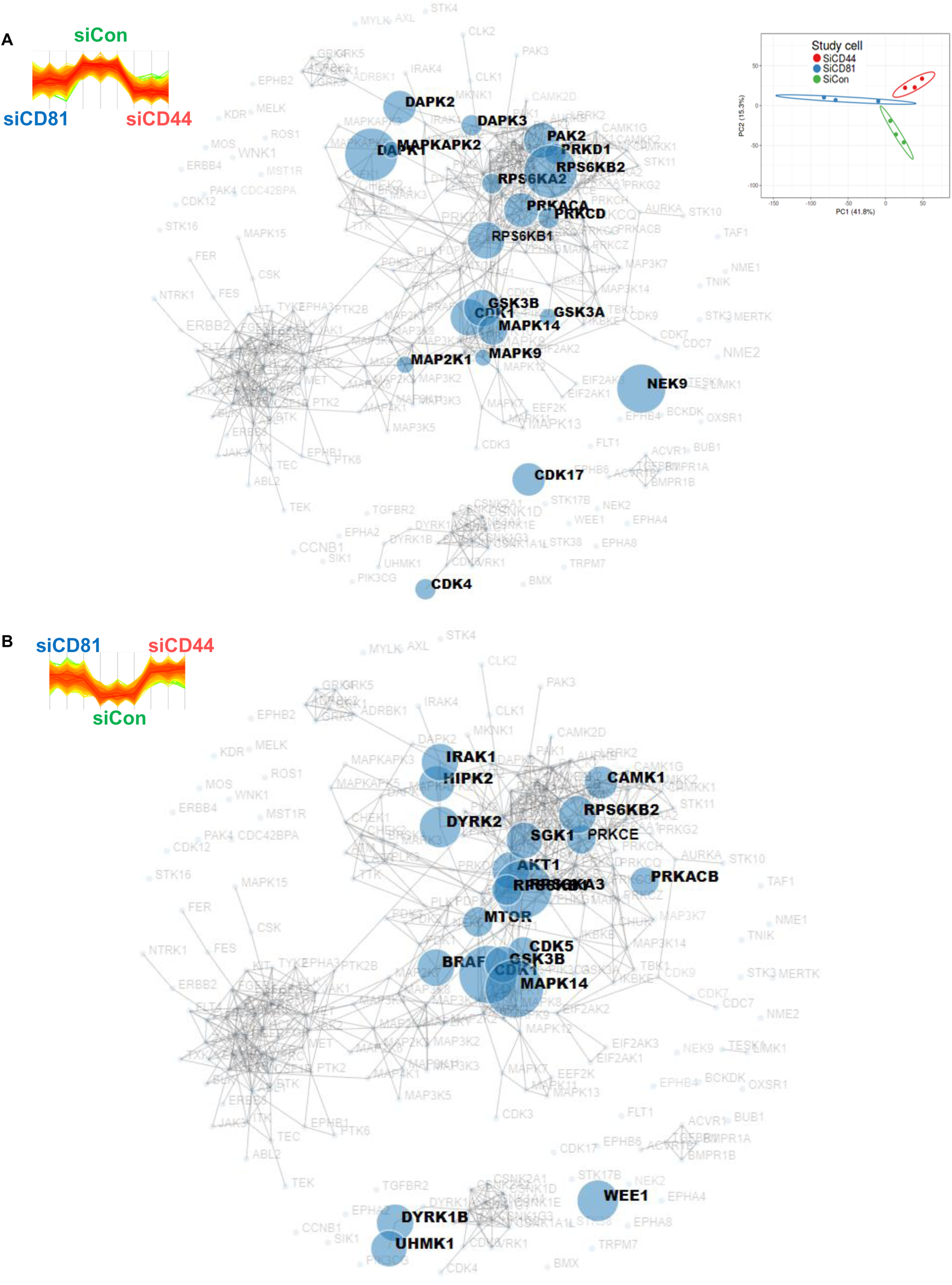
The kinase reactome networks (related to Fig. 2). The MaxQuant kinase reactome networks based on the downregulated (A) and upregulated (B) phosphorylation sites in both siCD81 and siCD44-transfected cells and known curation of kinase-substrate interactions in the literature.

**Suppl. Figure S5.**
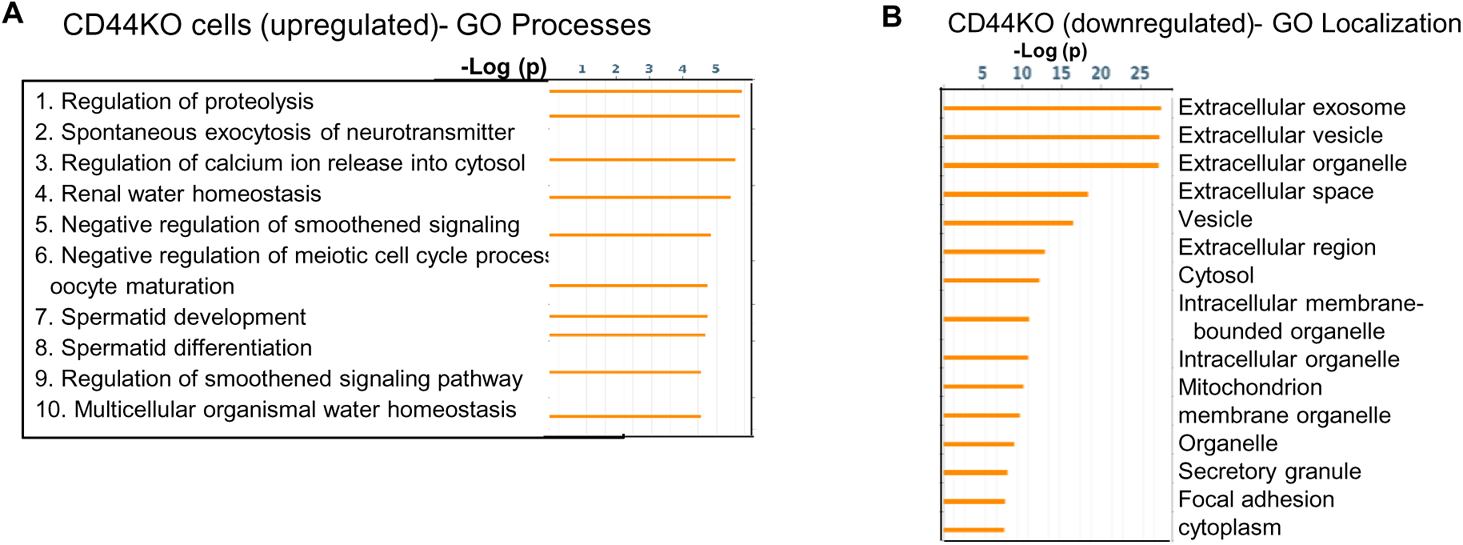
CD44 target pathways (related to Fig. 2) **A.** GO Localization analysis of downregulated proteins in pooled CD44KO cells compared to WT MDA-MB-231 cells (N=3 replicates, P<0.05). **B.** GO Processes analyses of global mass spectrometry-based upregulated proteins in the CD44KO cells in comparison to CD44 WT MDA-MB-231 cells.

**Suppl. Fig. S6.**
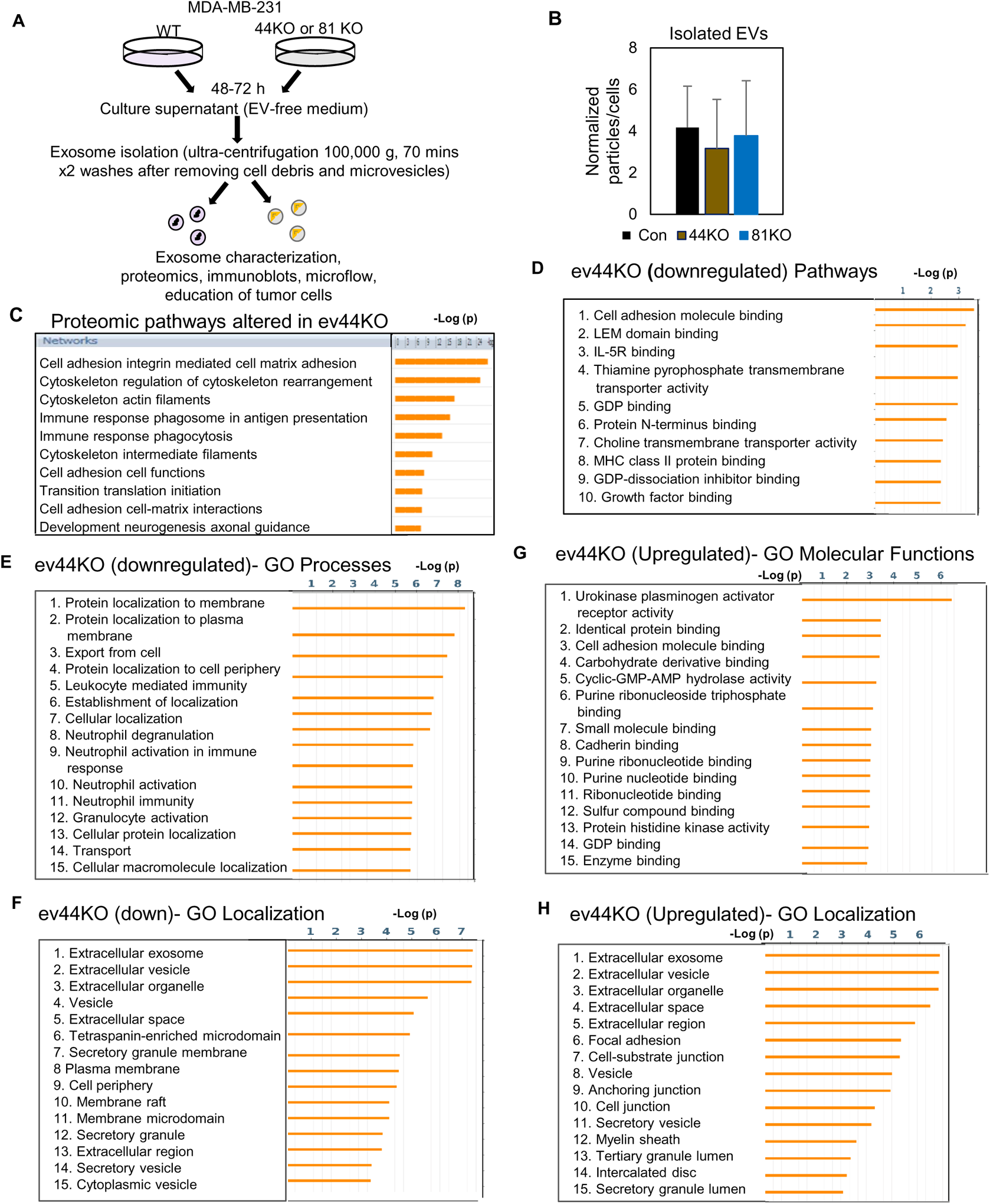
CD81 is required for exosome-induced effects on mammosphere formation (related to Figs. 1, 3). **A.** Schematic of EV isolation by ultracentrifugation steps and characterization. **B.** NTA analyses for purified EV particles per cell from WT, CD44KO and CD81KO MDA-MB-231 cells. No significant difference. Repeated 3 times. **C.** Proteomic pathways altered in the EVs derived from CD44KO cells (ev44KO) compared to evWT from WT cells. **D-H.** GO analyses of proteins downregulated or upregulated in the ev44KO versus evWT..

**Suppl. Fig. S7.**
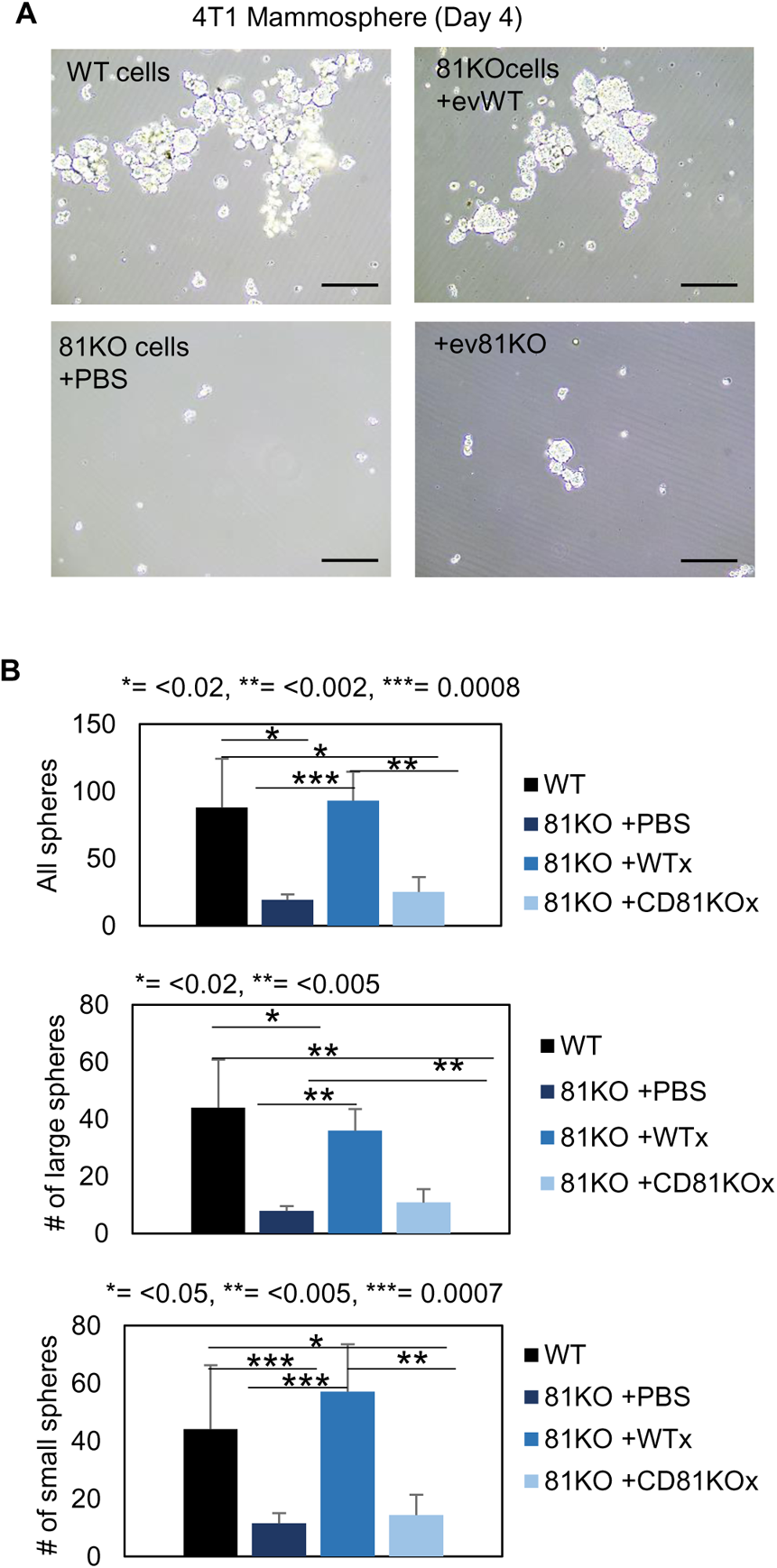
CD81 is required for exosome-induced effects on mammosphere formation (related to Figs. 1, 3). **A-B.** Mammosphere assessment of 4T1 cells, including WT cells and Cd81KO cells, the latter of which were educated with PBS or exosomes (10 µg for 1 week). 2,000 cells were seeded in 6 cm plates in mammosphere media, and the images were captured on day 4. Two-tailed student T-test was used.

**Suppl. Fig. S8.**
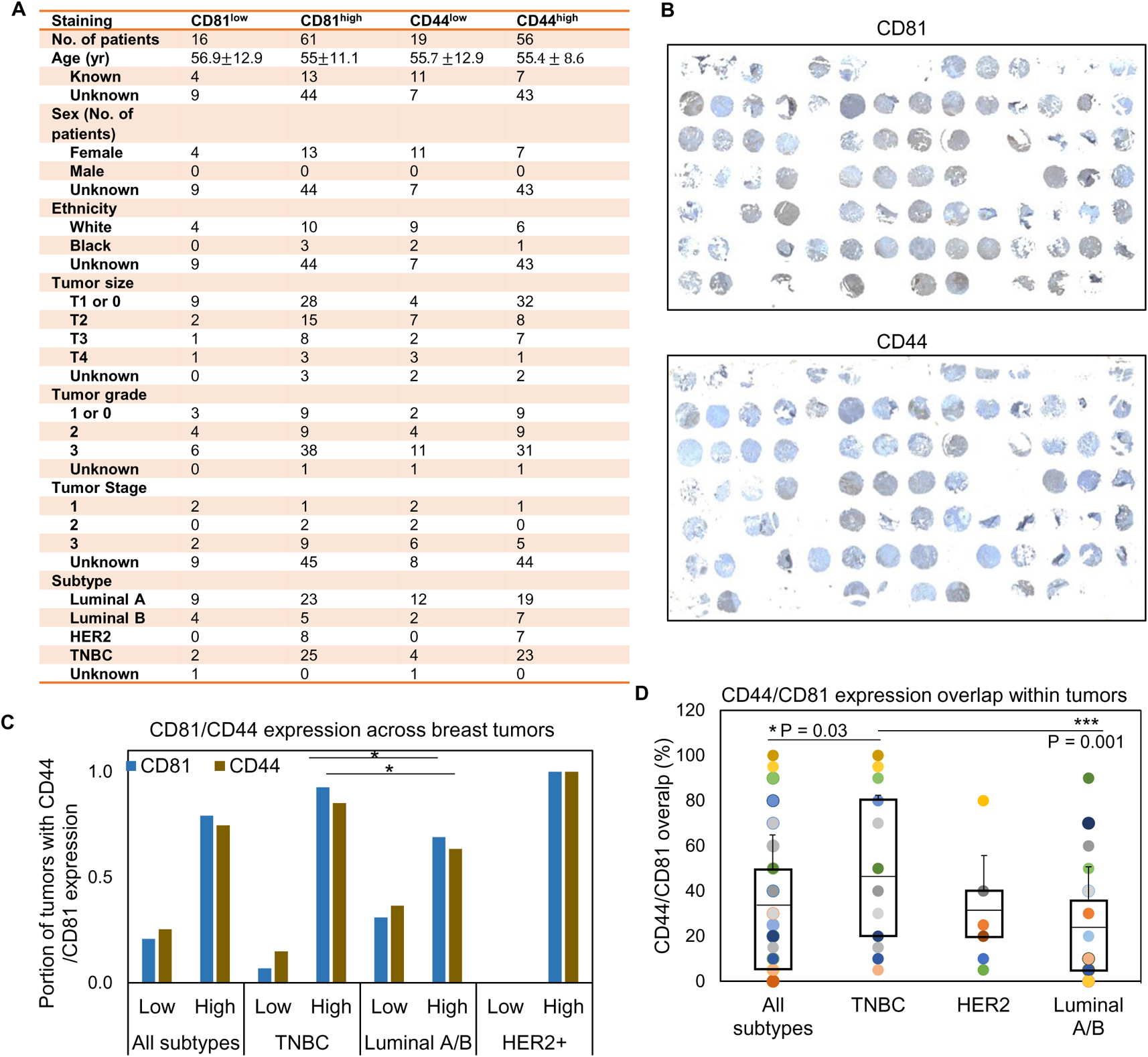
TMA clinical characteristics (related to Fig. 4) **A.** Table of tumor TMA breast cancer patient characteristics. **B.** Immunohistochemical staining of CD81 and CD44 expression (in brown) and hemotoxylin (in blue) in tissue microarray (TMA) of breast tumors (N=77 with CD81 IHC and N=75 with CD44 IHC). **C.** CD81 and CD44 expression levels were scored 1, 2, 3, or 4 ranging from very low, low, high, or very high. Very low and low were grouped into the low group and high and very high were grouped into the high group. Expression of CD81 and CD44 was assessed in all subtypes (N=77, N=75, respectively), TNBC (N=27, N=27, respectively), Luminal A/B (N=42, N=41, respectively), and HER2+ (N=8, N=7, respectively) patient tumors. Two-tailed student T-test * P < 0.05. **D.** Percentage (%) of tumor regions with overlapped CD81 and CD44 expression within each tumor (N=75 with both CD81 and CD44 IHC). Two-tailed student T-test P=0.03 and 0.001 showing TNBC with a higher overlap between CD81/CD44 than the average level of all subtypes as well as that of the Luminal A/B subtype.

**Suppl. Fig. S9.**
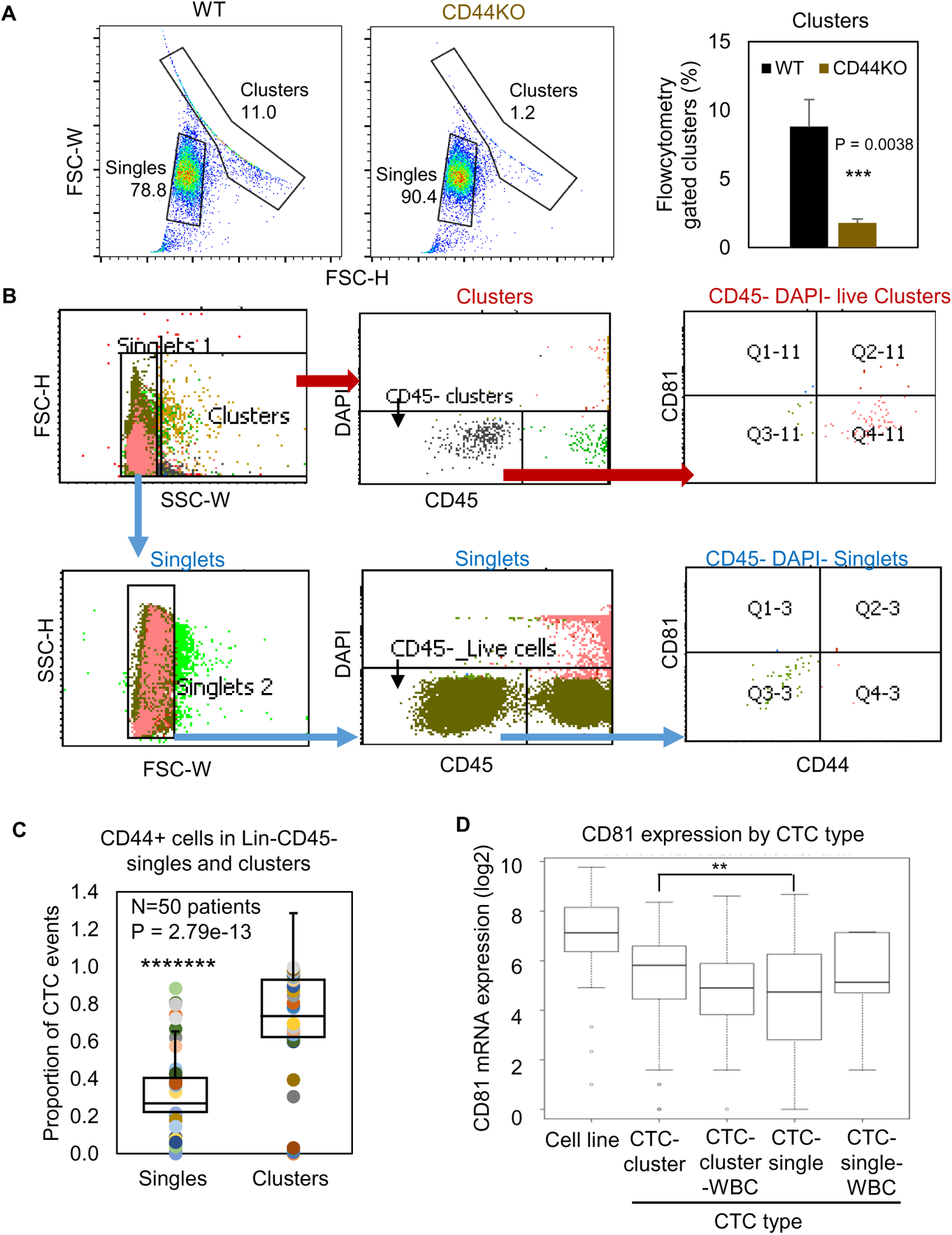
CTC analysis by flow cytometry and RNA sequencing (related to Fig. 4) **A.** Forward scatter channel (FSC)-gated singles and clusters of WT and CD44KO MDA-MB-231 cells in suspension (n=3 replicates). T test P=0.0001. **B.** Gating strategies for patient blood-isolated CD45^−^ single cells and clusters for CD81 and CD44 analyses. **C.** Plots of proportion of putative CD44+ CTC events (EpCAM^+/−^ clusters and singles in the blood of 50 patients with metastatic breast cancer, analyzed on flow cytometer). Two-tailed student T-test was used. **D.** mRNA expression of CD81 in single/clusters of CTCs from a publicly available dataset (ref 33 and 35). Wilcoxon p for CTC-single vs CTC-cluster = 0.0034.

**Suppl. Fig. S10.**
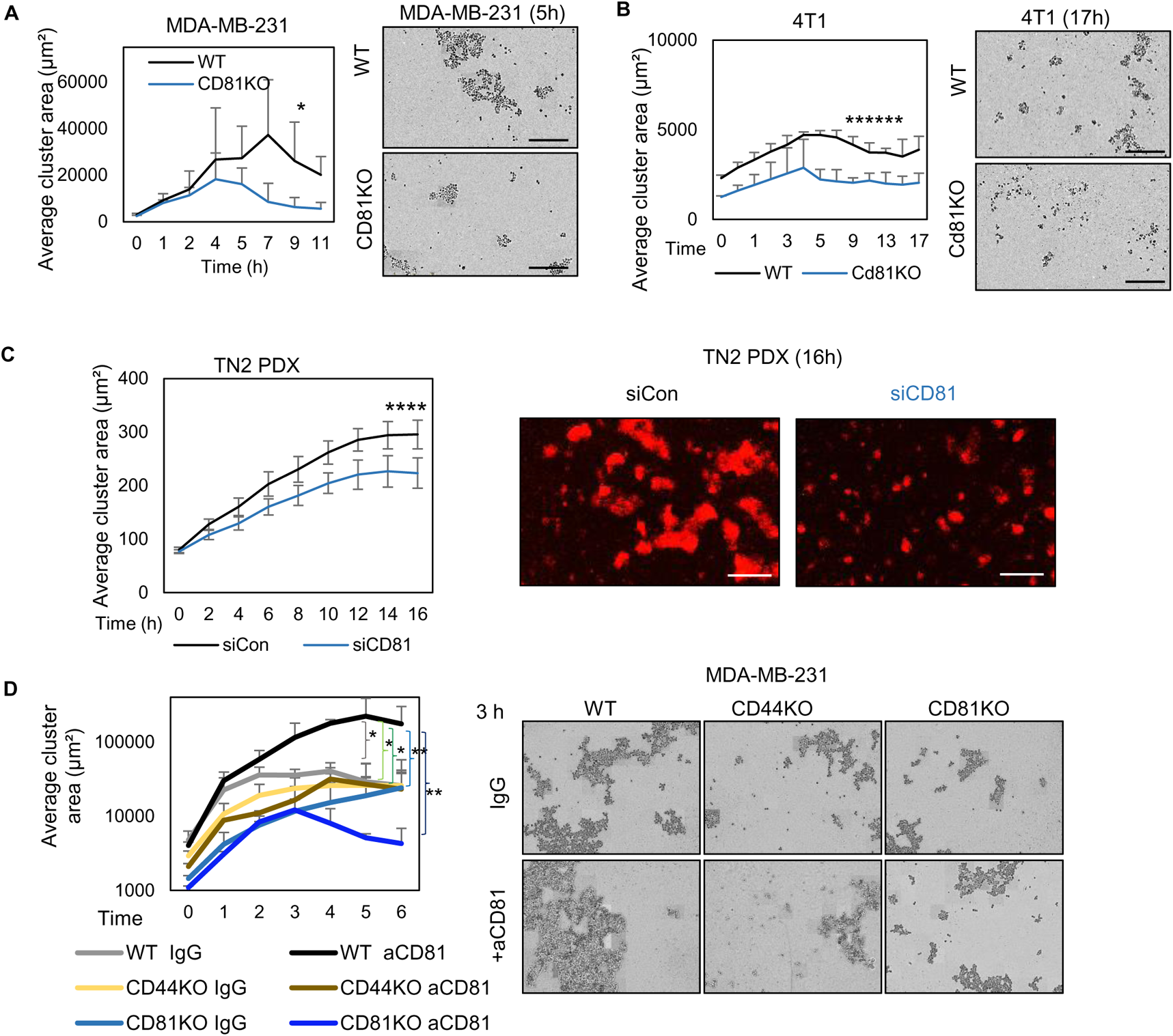
Tumor cells clustering (related to Fig. 4) **A-C.** IncuCyte images (right) and quantified tumor cell aggregation curves (left) of MDA-MB-231 WT/CD81KO cells (* P = 0.02) (A), 4T1 WT/Cd81KO cells (****** P <5.8E-08) (B), and TN2 PDX siCon/siCD81 (**** P = 0.0004) (C). Scale bar = 300 µm. Two-tailed student T-test was used. Repeated at least twice. **D.** Clustering analysis of MDA-MB-231 WT/CD44KO/CD81KO cells treated with 10 µg/ul IgG or anti-CD81 activating antibody. Two-tailed student T-test was used. P value comparing WT aCD81 to WT IgG *P = 0.022, to CD44KO IgG * P = 0.013, to CD44KO aCD81 * P = 0.011, to CD81KO IgG ** P = 0.008, and to CD81KO aCD81 ** P = 0.006.

**Suppl. Fig. S11.**
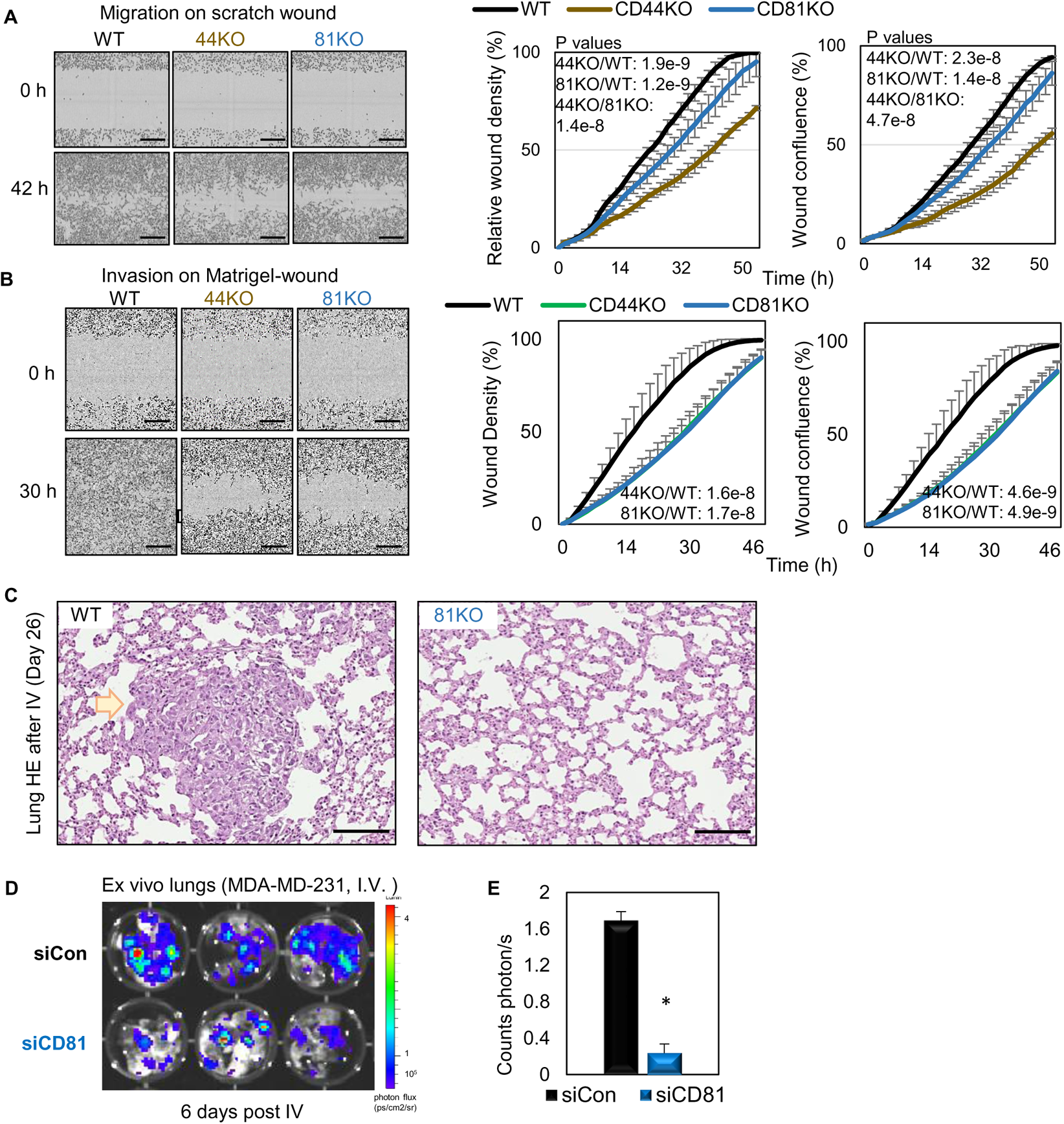
CD81 KD inhibits breast cancer cell metastasis (related to Figs. 6) **A.** Cell migration of MDA-MB-231 WT, CD44KO, and CD81KO cells to close scratch wound, analyzed by Incucyte time-lapse imaging in every 2 h. CD44KO and CD81KO groups took longer time to fill the wound gap. Two-tailed student T-test was used. Repeated twice. **B.** Cell invasion of MDA-MB-231 WT, CD44KO, and CD81KO cells on Matrigel-covered scratch wound was evaluated by Incucyte time-lapse imaging software which tracks wound closure every 2 h. After 30 h the control cell wounds began to close while the CD44KO and CD81KO groups took longer. Two-tailed student T-test was used. **C.** HE images of lung metastasis colonies (arrow pointed region) observed in the mice injected with WT cells and absent with 81KO MDA-MB-231 cells, harvested on day 26 after tail vein injection. Scale bar = 200 µm. **D, E.** CD81 control and knockdown MDA-MB-231 cells were evaluated for their metastatic potential. 100K cells were injected into mice by tail vein injection to observe lung colonization. 6 days post tail vein injection lung colonization was observed lower levels in the CD81KD group compared to the control. (n=3) (One-tailed student T-test * P = 0.04)

**Suppl Fig. S12.**
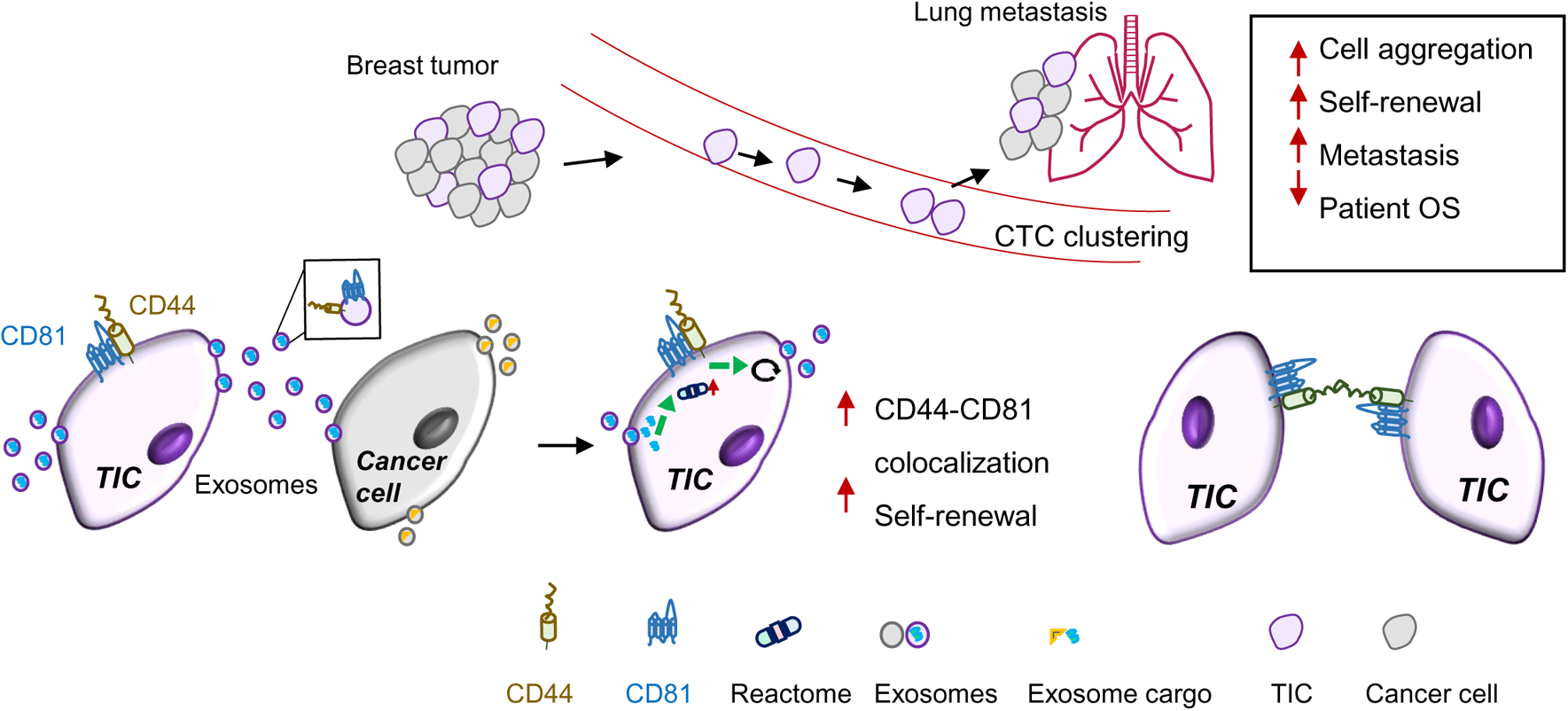
Schematic summary. Schematic summary of CD81 in interacting with CD44 on the cytoplasmic membrane of tumor-initiating cells (TIC), facilitating the exosome cargo packaging with CD44 and CD81, promoting exosome-induced self-renewal in recipient cells via phosphoreactome pathways, and strengthening CD44-mediated CTC cluster formation and lung metastasis.

## References

1. Sung H, Ferlay J, Siegel RL, Laversanne M, Soerjomataram I, Jemal A, Bray F. Global Cancer Statistics 2020: GLOBOCAN Estimates of Incidence and Mortality Worldwide for 36 Cancers in 185 Countries. CA Cancer J Clin. 2021;71(3):209–49. Epub 2021/02/05. doi: 10.3322/caac.21660. PubMed PMID: 33538338.

2. Perou CM, Sorlie T, Eisen MB, van de Rijn M, Jeffrey SS, Rees CA, Pollack JR, Ross DT, Johnsen H, Akslen LA, Fluge O, Pergamenschikov A, Williams C, Zhu SX, Lonning PE, Borresen-Dale AL, Brown PO, Botstein D. Molecular portraits of human breast tumours. Nature. 2000;406(6797):747–52. PubMed PMID: 10963602.

3. Sorlie T, Perou CM, Tibshirani R, Aas T, Geisler S, Johnsen H, Hastie T, Eisen MB, van de Rijn M, Jeffrey SS, Thorsen T, Quist H, Matese JC, Brown PO, Botstein D, Eystein Lonning P, Borresen-Dale AL. Gene expression patterns of breast carcinomas distinguish tumor subclasses with clinical implications. Proc Natl Acad Sci U S A. 2001;98(19):10869–74. PubMed PMID: 11553815.

4. Rakha EA, El-Sayed ME, Green AR, Lee AH, Robertson JF, Ellis IO. Prognostic markers in triple-negative breast cancer. Cancer. 2007;109(1):25–32. doi: 10.1002/cncr.22381. PubMed PMID: 17146782.

5. Haffty BG, Yang Q, Reiss M, Kearney T, Higgins SA, Weidhaas J, Harris L, Hait W, Toppmeyer D. Locoregional relapse and distant metastasis in conservatively managed triple negative early-stage breast cancer. J Clin Oncol. 2006;24(36):5652–7. doi: 10.1200/JCO.2006.06.5664. PubMed PMID: 17116942.

6. Foulkes WD, Smith IE, Reis-Filho JS. Triple-negative breast cancer. N Engl J Med. 2010;363(20):1938–48. Epub 2010/11/12. doi: 10.1056/NEJMra1001389. PubMed PMID: 21067385.

7. Dent R, Trudeau M, Pritchard KI, Hanna WM, Kahn HK, Sawka CA, Lickley LA, Rawlinson E, Sun P, Narod SA. Triple-negative breast cancer: clinical features and patterns of recurrence. Clin Cancer Res. 2007;13(15 Pt 1):4429–34. doi: 10.1158/1078-0432.CCR-06-3045. PubMed PMID: 17671126.

8. Diehn M, Cho RW, Lobo NA, Kalisky T, Dorie MJ, Kulp AN, Qian D, Lam JS, Ailles LE, Wong M, Joshua B, Kaplan MJ, Wapnir I, Dirbas FM, Somlo G, Garberoglio C, Paz B, Shen J, Lau SK, Quake SR, Brown JM, Weissman IL, Clarke MF. Association of reactive oxygen species levels and radioresistance in cancer stem cells. Nature. 2009;458(7239):780–3. doi: 10.1038/nature07733. PubMed PMID: 19194462; PMCID: PMC2778612.

9. Li X, Lewis MT, Huang J, Gutierrez C, Osborne CK, Wu MF, Hilsenbeck SG, Pavlick A, Zhang X, Chamness GC, Wong H, Rosen J, Chang JC. Intrinsic resistance of tumorigenic breast cancer cells to chemotherapy. J Natl Cancer Inst. 2008;100(9):672–9. Epub 2008/05/01. doi: djn123 [pii] 10.1093/jnci/djn123. PubMed PMID: 18445819.

10. Al-Hajj M, Wicha MS, Benito-Hernandez A, Morrison SJ, Clarke MF. Prospective identification of tumorigenic breast cancer cells. Proc Natl Acad Sci U S A. 2003;100(7):3983–8. Epub 2003/03/12. doi: 10.1073/pnas.0530291100 0530291100 [pii]. PubMed PMID: 12629218; PMCID: 153034.

11. Charafe-Jauffret E, Ginestier C, Iovino F, Tarpin C, Diebel M, Esterni B, Houvenaeghel G, Extra JM, Bertucci F, Jacquemier J, Xerri L, Dontu G, Stassi G, Xiao Y, Barsky SH, Birnbaum D, Viens P, Wicha MS. Aldehyde dehydrogenase 1-positive cancer stem cells mediate metastasis and poor clinical outcome in inflammatory breast cancer. Clinical cancer research : an official journal of the American Association for Cancer Research. 2010;16(1):45–55. doi: 10.1158/1078-0432.CCR-09-1630. PubMed PMID: 20028757; PMCID: 2874875.

12. Giordano A, Gao H, Cohen EN, Anfossi S, Khoury J, Hess K, Krishnamurthy S, Tin S, Cristofanilli M, Hortobagyi GN, Woodward WA, Lucci A, Reuben JM. Clinical relevance of cancer stem cells in bone marrow of early breast cancer patients. Annals of oncology : official journal of the European Society for Medical Oncology / ESMO. 2013;24(10):2515–21. doi: 10.1093/annonc/mdt223. PubMed PMID: 23798614; PMCID: 3784333.

13. Idowu MO, Kmieciak M, Dumur C, Burton RS, Grimes MM, Powers CN, Manjili MH. CD44(+)/CD24(-/low) cancer stem/progenitor cells are more abundant in triple-negative invasive breast carcinoma phenotype and are associated with poor outcome. Human pathology. 2012;43(3):364–73. doi: 10.1016/j.humpath.2011.05.005. PubMed PMID: 21835433.

14. Lin Y, Zhong Y, Guan H, Zhang X, Sun Q. CD44+/CD24- phenotype contributes to malignant relapse following surgical resection and chemotherapy in patients with invasive ductal carcinoma. Journal of experimental & clinical cancer research : CR. 2012;31:59. doi: 10.1186/1756-9966-31-59. PubMed PMID: 22762532; PMCID: 3432011.

15. Owens TW, Naylor MJ. Breast cancer stem cells. Frontiers in physiology. 2013;4:225. doi: 10.3389/fphys.2013.00225. PubMed PMID: 23986719; PMCID: 3753536.

16. Liu H, Patel MR, Prescher JA, Patsialou A, Qian D, Lin J, Wen S, Chang YF, Bachmann MH, Shimono Y, Dalerba P, Adorno M, Lobo N, Bueno J, Dirbas FM, Goswami S, Somlo G, Condeelis J, Contag CH, Gambhir SS, Clarke MF. Cancer stem cells from human breast tumors are involved in spontaneous metastases in orthotopic mouse models. Proc Natl Acad Sci U S A. 2010;107(42):18115–20. Epub 2010/10/06. doi: 1006732107 [pii] 10.1073/pnas.1006732107. PubMed PMID: 20921380.

17. Liu S, Wicha MS. Targeting breast cancer stem cells. Journal of clinical oncology : official journal of the American Society of Clinical Oncology. 2010;28(25):4006–12. doi: 10.1200/JCO.2009.27.5388. PubMed PMID: 20498387.

18. Korkaya H, Wicha MS. HER-2, notch, and breast cancer stem cells: targeting an axis of evil. Clinical cancer research : an official journal of the American Association for Cancer Research. 2009;15(6):1845–7. doi: 10.1158/1078-0432.CCR-08-3087. PubMed PMID: 19276254.

19. Adorno-Cruz V, Kibria G, Liu X, Doherty M, Junk DJ, Guan D, Hubert C, Venere M, Mulkearns-Hubert E, Sinyuk M, Alvarado A, Caplan AI, Rich J, Gerson SL, Lathia J, Liu H. Cancer stem cells: targeting the roots of cancer, seeds of metastasis, and sources of therapy resistance. Cancer Res. 2015;75(6):924–9. doi: 10.1158/0008-5472.CAN-14-3225. PubMed PMID: 25604264; PMCID: PMC4359955.

20. Ramos EK, Hoffmann AD, Gerson SL, Liu H. New Opportunities and Challenges to Defeat Cancer Stem Cells. Trends Cancer. 2017;3(11):780–96. doi: 10.1016/j.trecan.2017.08.007. PubMed PMID: 29120754.

21. Bhola NE, Balko JM, Dugger TC, Kuba MG, Sanchez V, Sanders M, Stanford J, Cook RS, Arteaga CL. TGF-beta inhibition enhances chemotherapy action against triple-negative breast cancer. J Clin Invest. 2013;123(3):1348–58. doi: 10.1172/Jci65416. PubMed PMID: WOS:000315749400044.

22. Calcagno AM, Salcido CD, Gillet JP, Wu CP, Fostel JM, Mumau MD, Gottesman MM, Varticovski L, Ambudkar SV. Prolonged Drug Selection of Breast Cancer Cells and Enrichment of Cancer Stem Cell Characteristics. J Natl Cancer I. 2010;102(21):1637–52. doi: 10.1093/jnci/djq361. PubMed PMID: WOS:000283923300009.

23. Charafe-Jauffret E, Ginestier C, Iovino F, Tarpin C, Diebel M, Esterni B, Houvenaeghel G, Extra JM, Bertucci F, Jacquemier J, Xerri L, Dontu G, Stassi G, Xiao Y, Barsky SH, Birnbaum D, Viens P, Wicha MS. Aldehyde Dehydrogenase 1-Positive Cancer Stem Cells Mediate Metastasis and Poor Clinical Outcome in Inflammatory Breast Cancer. Clin Cancer Res. 2010;16(1):45–55. doi: 10.1158/1078-0432.Ccr-09-1630. PubMed PMID: WOS:000278404500006.

24. Idowu MO, Kmieciak M, Dumur C, Burton RS, Grimes MM, Powers CN, Manjili MH. CD44(+)/CD24(-/low) cancer stem/progenitor cells are more abundant in triple-negative invasive breast carcinoma phenotype and are associated with poor outcome. Hum Pathol. 2012;43(3):364–73. doi: 10.1016/j.humpath.2011.05.005. PubMed PMID: WOS:000300590800008.

25. Li XX, Lewis MT, Huang J, Gutierrez C, Osborne CK, Wu MF, Hilsenbeck SG, Pavlick A, Zhang XM, Chamness GC, Wong H, Rosen J, Chang JC. Intrinsic resistance of tumorigenic breast cancer cells to chemotherapy. J Natl Cancer I. 2008;100(9):672–9. doi: 10.1093/jnci/djn123. PubMed PMID: WOS:000255757200015.

26. Lin Y, Zhong Y, Guan H, Zhang XH, Sun Q. CD44(+)/CD24(-) phenotype contributes to malignant relapse following surgical resection and chemotherapy in patients with invasive ductal carcinoma. J Exp Clin Canc Res. 2012;31. doi: Artn 59 10.1186/1756-9966-31-59. PubMed PMID: WOS:000308438800001.

27. Owens TW, Naylor MJ. Breast cancer stem cells. Front Physiol. 2013;4. doi: ARTN 225 10.3389/fphys.2013.00225. PubMed PMID: WOS:000346774000222.

28. Phillips TM, McBride WH, Pajonk F. The response of CD24(-/low)/CD44(+) breast cancer-initiating cells to radiation. J Natl Cancer I. 2006;98(24):1777–85. doi: 10.1093/jnci/djj495. PubMed PMID: WOS:000242973400009.

29. Creighton CJ, Li XX, Landis M, Dixon JM, Neumeister VM, Sjolund A, Rimm DL, Wong H, Rodriguez A, Herschkowitz JI, Fan C, Zhang XM, He XP, Pavlick A, Gutierrez MC, Renshaw L, Larionov AA, Faratian D, Hilsenbeck SG, Perou CM, Lewis MT, Rosen JM, Chang JC. Residual breast cancers after conventional therapy display mesenchymal as well as tumor-initiating features. P Natl Acad Sci USA. 2009;106(33):13820–5. doi: 10.1073/pnas.0905718106. PubMed PMID: WOS:000269078700037.

30. Shats I, Gatza ML, Chang JT, Mori S, Wang JL, Rich J, Nevins JR. Using a Stem Cell-Based Signature to Guide Therapeutic Selection in Cancer. Cancer Res. 2011;71(5):1772–80. doi: 10.1158/0008-5472.Can-10-1735. PubMed PMID: WOS:000287845300028.

31. Lathia J, Liu H, Matei D. The Clinical Impact of Cancer Stem Cells. Oncologist. 2020;25(2):123–31. Epub 2020/02/12. doi: 10.1634/theoncologist.2019-0517. PubMed PMID: 32043793; PMCID: PMC7011672.

32. Fernandez SV, Bingham C, Fittipaldi P, Austin L, Palazzo J, Palmer G, Alpaugh K, Cristofanilli M. TP53 mutations detected in circulating tumor cells present in the blood of metastatic triple negative breast cancer patients. Breast Cancer Res. 2014;16(5):445. Epub 2014/10/14. doi: 10.1186/s13058-014-0445-3. PubMed PMID: 25307991; PMCID: PMC4303125.

33. Gkountela S, Castro-Giner F, Szczerba BM, Vetter M, Landin J, Scherrer R, Krol I, Scheidmann MC, Beisel C, Stirnimann CU, Kurzeder C, Heinzelmann-Schwarz V, Rochlitz C, Weber WP, Aceto N. Circulating Tumor Cell Clustering Shapes DNA Methylation to Enable Metastasis Seeding. Cell. 2019;176(1-2):98–112 e14. Epub 2019/01/12. doi: 10.1016/j.cell.2018.11.046. PubMed PMID: 30633912; PMCID: PMC6363966.

34. Mu Z, Wang C, Ye Z, Austin L, Civan J, Hyslop T, Palazzo JP, Jaslow R, Li B, Myers RE, Jiang J, Xing J, Yang H, Cristofanilli M. Prospective assessment of the prognostic value of circulating tumor cells and their clusters in patients with advanced-stage breast cancer. Breast Cancer Res Treat. 2015;154(3):563–71. doi: 10.1007/s10549-015-3636-4. PubMed PMID: 26573830.

35. Aceto N, Bardia A, Miyamoto DT, Donaldson MC, Wittner BS, Spencer JA, Yu M, Pely A, Engstrom A, Zhu H, Brannigan BW, Kapur R, Stott SL, Shioda T, Ramaswamy S, Ting DT, Lin CP, Toner M, Haber DA, Maheswaran S. Circulating tumor cell clusters are oligoclonal precursors of breast cancer metastasis. Cell. 2014;158(5):1110–22. Epub 2014/08/30. doi: 10.1016/j.cell.2014.07.013. PubMed PMID: 25171411; PMCID: PMC4149753.

36. Liu X, Taftaf R, Kawaguchi M, Chang YF, Chen W, Entenberg D, Zhang Y, Gerratana L, Huang S, Patel DB, Tsui E, Adorno-Cruz V, Chirieleison SM, Cao Y, Harney AS, Patel S, Patsialou A, Shen Y, Avril S, Gilmore HL, Lathia JD, Abbott DW, Cristofanilli M, Condeelis JS, Liu H. Homophilic CD44 Interactions Mediate Tumor Cell Aggregation and Polyclonal Metastasis in Patient-Derived Breast Cancer Models. Cancer Discov. 2019;9(1):96–113. Epub 2018/10/27. doi: 10.1158/2159-8290.CD-18-0065. PubMed PMID: 30361447; PMCID: PMC6328322.

37. Ting DT, Wittner BS, Ligorio M, Vincent Jordan N, Shah AM, Miyamoto DT, Aceto N, Bersani F, Brannigan BW, Xega K, Ciciliano JC, Zhu H, MacKenzie OC, Trautwein J, Arora KS, Shahid M, Ellis HL, Qu N, Bardeesy N, Rivera MN, Deshpande V, Ferrone CR, Kapur R, Ramaswamy S, Shioda T, Toner M, Maheswaran S, Haber DA. Single-cell RNA sequencing identifies extracellular matrix gene expression by pancreatic circulating tumor cells. Cell Rep. 2014;8(6):1905–18. Epub 2014/09/23. doi: 10.1016/j.celrep.2014.08.029. PubMed PMID: 25242334; PMCID: PMC4230325.

38. McAndrews KM, Che SPY, LeBleu VS, Kalluri R. Effective delivery of STING agonist using exosomes suppresses tumor growth and enhances anti-tumor immunity. J Biol Chem. 2021:100523. Epub 2021/03/13. doi: 10.1016/j.jbc.2021.100523. PubMed PMID: 33711340; PMCID: PMC8042450.

39. Kahlert C, Kalluri R. Exosomes in tumor microenvironment influence cancer progression and metastasis. J Mol Med (Berl). 2013;91(4):431–7. doi: 10.1007/s00109-013-1020-6. PubMed PMID: 23519402; PMCID: PMC4073669.

40. Valadi H, Ekstrom K, Bossios A, Sjostrand M, Lee JJ, Lotvall JO. Exosome-mediated transfer of mRNAs and microRNAs is a novel mechanism of genetic exchange between cells. Nat Cell Biol. 2007;9(6):654–9. doi: 10.1038/ncb1596. PubMed PMID: 17486113.

41. Peinado H, Aleckovic M, Lavotshkin S, Matei I, Costa-Silva B, Moreno-Bueno G, Hergueta-Redondo M, Williams C, Garcia-Santos G, Ghajar C, Nitadori-Hoshino A, Hoffman C, Badal K, Garcia BA, Callahan MK, Yuan J, Martins VR, Skog J, Kaplan RN, Brady MS, Wolchok JD, Chapman PB, Kang Y, Bromberg J, Lyden D. Melanoma exosomes educate bone marrow progenitor cells toward a pro-metastatic phenotype through MET. Nat Med. 2012;18(6):883–91. Epub 2012/05/29. doi: 10.1038/nm.2753. PubMed PMID: 22635005; PMCID: PMC3645291.

42. M C. Exosome-based candidates move into the clinic. Nat Rev Drug Discovery. 2020;2021.

43. Hoshino A, Costa-Silva B, Shen TL, Rodrigues G, Hashimoto A, Tesic Mark M, Molina H, Kohsaka S, Di Giannatale A, Ceder S, Singh S, Williams C, Soplop N, Uryu K, Pharmer L, King T, Bojmar L, Davies AE, Ararso Y, Zhang T, Zhang H, Hernandez J, Weiss JM, Dumont-Cole VD, Kramer K, Wexler LH, Narendran A, Schwartz GK, Healey JH, Sandstrom P, Labori KJ, Kure EH, Grandgenett PM, Hollingsworth MA, de Sousa M, Kaur S, Jain M, Mallya K, Batra SK, Jarnagin WR, Brady MS, Fodstad O, Muller V, Pantel K, Minn AJ, Bissell MJ, Garcia BA, Kang Y, Rajasekhar VK, Ghajar CM, Matei I, Peinado H, Bromberg J, Lyden D. Tumour exosome integrins determine organotropic metastasis. Nature. 2015;527(7578):329–35. doi: 10.1038/nature15756. PubMed PMID: 26524530; PMCID: PMC4788391.

44. Kamerkar S, LeBleu VS, Sugimoto H, Yang S, Ruivo CF, Melo SA, Lee JJ, Kalluri R. Exosomes facilitate therapeutic targeting of oncogenic KRAS in pancreatic cancer. Nature. 2017;546(7659):498–503. doi: 10.1038/nature22341. PubMed PMID: 28607485; PMCID: PMC5538883.

45. Kibria G, Ramos EK, Wan Y, Gius DR, Liu H. Exosomes as a Drug Delivery System in Cancer Therapy: Potential and Challenges. Mol Pharm. 2018;15(9):3625–33. doi: 10.1021/acs.molpharmaceut.8b00277. PubMed PMID: 29771531.

46. Bobrie A, Colombo M, Raposo G, Thery C. Exosome secretion: molecular mechanisms and roles in immune responses. Traffic. 2011;12(12):1659–68. doi: 10.1111/j.1600-0854.2011.01225.x. PubMed PMID: 21645191.

47. Thery C, Amigorena S, Raposo G, Clayton A. Isolation and characterization of exosomes from cell culture supernatants and biological fluids. Curr Protoc Cell Biol. 2006;Chapter 3:Unit 3 22. doi: 10.1002/0471143030.cb0322s30. PubMed PMID: 18228490.

48. Iero M, Valenti R, Huber V, Filipazzi P, Parmiani G, Fais S, Rivoltini L. Tumour-released exosomes and their implications in cancer immunity. Cell Death Differ. 2008;15(1):80–8. doi: 10.1038/sj.cdd.4402237. PubMed PMID: WOS:000251820900010.

49. Kahlert C, Kalluri R. Exosomes in tumor microenvironment influence cancer progression and metastasis. J Mol Med. 2013;91(4):431–7. doi: 10.1007/s00109-013-1020-6. PubMed PMID: WOS:000317077200003.

50. Hoshino A, Costa-Silva B, Shen TL, Rodrigues G, Hashimoto A, Mark MT, Molina H, Kohsaka S, Di Giannatale A, Ceder S, Singh S, Williams C, Soplop N, Uryu K, Pharmer L, King T, Bojmar L, Davies AE, Ararso Y, Zhang T, Zhang H, Hernandez J, Weiss JM, Dumont-Cole VD, Kramer K, Wexler LH, Narendran A, Schwartz GK, Healey JH, Sandstrom P, Labori KJ, Kure EH, Grandgenett PM, Hollingsworth MA, de Sousa M, Kaur S, Jain M, Mallya K, Batra SK, Jarnagin WR, Brady MS, Fodstad O, Muller V, Pantel K, Minn AJ, Bissell MJ, Garcia BA, Kang Y, Rajasekhar VK, Ghajar CM, Matei I, Peinado H, Bromberg J, Lyden D. Tumour exosome integrins determine organotropic metastasis. Nature. 2015;527(7578):329-+. doi: 10.1038/nature15756. PubMed PMID: WOS:000365356800046.

51. Thery C, Witwer KW, Aikawa E, Alcaraz MJ, Anderson JD, Andriantsitohaina R, Antoniou A, Arab T, Archer F, Atkin-Smith GK, Ayre DC, Bach JM, Bachurski D, Baharvand H, Balaj L, Baldacchino S, Bauer NN, Baxter AA, Bebawy M, Beckham C, Bedina Zavec A, Benmoussa A, Berardi AC, Bergese P, Bielska E, Blenkiron C, Bobis-Wozowicz S, Boilard E, Boireau W, Bongiovanni A, Borras FE, Bosch S, Boulanger CM, Breakefield X, Breglio AM, Brennan MA, Brigstock DR, Brisson A, Broekman ML, Bromberg JF, Bryl-Gorecka P, Buch S, Buck AH, Burger D, Busatto S, Buschmann D, Bussolati B, Buzas EI, Byrd JB, Camussi G, Carter DR, Caruso S, Chamley LW, Chang YT, Chen C, Chen S, Cheng L, Chin AR, Clayton A, Clerici SP, Cocks A, Cocucci E, Coffey RJ, Cordeiro-da-Silva A, Couch Y, Coumans FA, Coyle B, Crescitelli R, Criado MF, D’Souza-Schorey C, Das S, Datta Chaudhuri A, de Candia P, De Santana EF, De Wever O, Del Portillo HA, Demaret T, Deville S, Devitt A, Dhondt B, Di Vizio D, Dieterich LC, Dolo V, Dominguez Rubio AP, Dominici M, Dourado MR, Driedonks TA, Duarte FV, Duncan HM, Eichenberger RM, Ekstrom K, El Andaloussi S, Elie-Caille C, Erdbrugger U, Falcon-Perez JM, Fatima F, Fish JE, Flores-Bellver M, Forsonits A, Frelet-Barrand A, Fricke F, Fuhrmann G, Gabrielsson S, Gamez-Valero A, Gardiner C, Gartner K, Gaudin R, Gho YS, Giebel B, Gilbert C, Gimona M, Giusti I, Goberdhan DC, Gorgens A, Gorski SM, Greening DW, Gross JC, Gualerzi A, Gupta GN, Gustafson D, Handberg A, Haraszti RA, Harrison P, Hegyesi H, Hendrix A, Hill AF, Hochberg FH, Hoffmann KF, Holder B, Holthofer H, Hosseinkhani B, Hu G, Huang Y, Huber V, Hunt S, Ibrahim AG, Ikezu T, Inal JM, Isin M, Ivanova A, Jackson HK, Jacobsen S, Jay SM, Jayachandran M, Jenster G, Jiang L, Johnson SM, Jones JC, Jong A, Jovanovic-Talisman T, Jung S, Kalluri R, Kano SI, Kaur S, Kawamura Y, Keller ET, Khamari D, Khomyakova E, Khvorova A, Kierulf P, Kim KP, Kislinger T, Klingeborn M, Klinke DJ, 2nd, Kornek M, Kosanovic MM, Kovacs AF, Kramer-Albers EM, Krasemann S, Krause M, Kurochkin IV, Kusuma GD, Kuypers S, Laitinen S, Langevin SM, Languino LR, Lannigan J, Lasser C, Laurent LC, Lavieu G, Lazaro-Ibanez E, Le Lay S, Lee MS, Lee YXF, Lemos DS, Lenassi M, Leszczynska A, Li IT, Liao K, Libregts SF, Ligeti E, Lim R, Lim SK, Line A, Linnemannstons K, Llorente A, Lombard CA, Lorenowicz MJ, Lorincz AM, Lotvall J, Lovett J, Lowry MC, Loyer X, Lu Q, Lukomska B, Lunavat TR, Maas SL, Malhi H, Marcilla A, Mariani J, Mariscal J, Martens-Uzunova ES, Martin-Jaular L, Martinez MC, Martins VR, Mathieu M, Mathivanan S, Maugeri M, McGinnis LK, McVey MJ, Meckes DG, Jr., Meehan KL, Mertens I, Minciacchi VR, Moller A, Moller Jorgensen M, Morales-Kastresana A, Morhayim J, Mullier F, Muraca M, Musante L, Mussack V, Muth DC, Myburgh KH, Najrana T, Nawaz M, Nazarenko I, Nejsum P, Neri C, Neri T, Nieuwland R, Nimrichter L, Nolan JP, Nolte-’t Hoen EN, Noren Hooten N, O’Driscoll L, O’Grady T, O’Loghlen A, Ochiya T, Olivier M, Ortiz A, Ortiz LA, Osteikoetxea X, Ostergaard O, Ostrowski M, Park J, Pegtel DM, Peinado H, Perut F, Pfaffl MW, Phinney DG, Pieters BC, Pink RC, Pisetsky DS, Pogge von Strandmann E, Polakovicova I, Poon IK, Powell BH, Prada I, Pulliam L, Quesenberry P, Radeghieri A, Raffai RL, Raimondo S, Rak J, Ramirez MI, Raposo G, Rayyan MS, Regev-Rudzki N, Ricklefs FL, Robbins PD, Roberts DD, Rodrigues SC, Rohde E, Rome S, Rouschop KM, Rughetti A, Russell AE, Saa P, Sahoo S, Salas-Huenuleo E, Sanchez C, Saugstad JA, Saul MJ, Schiffelers RM, Schneider R, Schoyen TH, Scott A, Shahaj E, Sharma S, Shatnyeva O, Shekari F, Shelke GV, Shetty AK, Shiba K, Siljander PR, Silva AM, Skowronek A, Snyder OL, 2nd, Soares RP, Sodar BW, Soekmadji C, Sotillo J, Stahl PD, Stoorvogel W, Stott SL, Strasser EF, Swift S, Tahara H, Tewari M, Timms K, Tiwari S, Tixeira R, Tkach M, Toh WS, Tomasini R, Torrecilhas AC, Tosar JP, Toxavidis V, Urbanelli L, Vader P, van Balkom BW, van der Grein SG, Van Deun J, van Herwijnen MJ, Van Keuren-Jensen K, van Niel G, van Royen ME, van Wijnen AJ, Vasconcelos MH, Vechetti IJ, Jr., Veit TD, Vella LJ, Velot E, Verweij FJ, Vestad B, Vinas JL, Visnovitz T, Vukman KV, Wahlgren J, Watson DC, Wauben MH, Weaver A, Webber JP, Weber V, Wehman AM, Weiss DJ, Welsh JA, Wendt S, Wheelock AM, Wiener Z, Witte L, Wolfram J, Xagorari A, Xander P, Xu J, Yan X, Yanez-Mo M, Yin H, Yuana Y, Zappulli V, Zarubova J, Zekas V, Zhang JY, Zhao Z, Zheng L, Zheutlin AR, Zickler AM, Zimmermann P, Zivkovic AM, Zocco D, Zuba-Surma EK. Minimal information for studies of extracellular vesicles 2018 (MISEV2018): a position statement of the International Society for Extracellular Vesicles and update of the MISEV2014 guidelines. J Extracell Vesicles. 2018;7(1):1535750. Epub 2019/01/15. doi: 10.1080/20013078.2018.1535750. PubMed PMID: 30637094; PMCID: PMC6322352.

52. Cao Y, Shen Y. Bayesian Active Learning for Optimization and Uncertainty Quantification in Protein Docking. J Chem Theory Comput. 2020;16(8):5334–47. Epub 2020/06/20. doi: 10.1021/acs.jctc.0c00476. PubMed PMID: 32558561; PMCID: PMC7429362.

53. Kawaguchi M, Dashzeveg N, Cao Y, Jia Y, Liu X, Shen Y, Liu H. Extracellular Domains I and II of cell-surface glycoprotein CD44 mediate its trans-homophilic dimerization and tumor cluster aggregation. J Biol Chem. 2020;295(9):2640–9. Epub 2020/01/24. doi: 10.1074/jbc.RA119.010252. PubMed PMID: 31969394; PMCID: PMC7049959.

54. Zimmerman B, Kelly B, McMillan BJ, Seegar TCM, Dror RO, Kruse AC, Blacklow SC. Crystal Structure of a Full-Length Human Tetraspanin Reveals a Cholesterol-Binding Pocket. Cell. 2016;167(4):1041–51 e11. Epub 2016/11/25. doi: 10.1016/j.cell.2016.09.056. PubMed PMID: 27881302; PMCID: PMC5127602.

55. Yang J, Yan R, Roy A, Xu D, Poisson J, Zhang Y. The I-TASSER Suite: protein structure and function prediction. Nat Methods. 2015;12(1):7–8. doi: 10.1038/nmeth.3213. PubMed PMID: 25549265; PMCID: PMC4428668.

56. Kozakov D, Hall DR, Xia B, Porter KA, Padhorny D, Yueh C, Beglov D, Vajda S. The ClusPro web server for protein-protein docking. Nat Protoc. 2017;12(2):255–78. doi: 10.1038/nprot.2016.169. PubMed PMID: 28079879; PMCID: PMC5540229.

57. Tripathi S, Pohl MO, Zhou Y, Rodriguez-Frandsen A, Wang G, Stein DA, Moulton HM, DeJesus P, Che J, Mulder LC, Yanguez E, Andenmatten D, Pache L, Manicassamy B, Albrecht RA, Gonzalez MG, Nguyen Q, Brass A, Elledge S, White M, Shapira S, Hacohen N, Karlas A, Meyer TF, Shales M, Gatorano A, Johnson JR, Jang G, Johnson T, Verschueren E, Sanders D, Krogan N, Shaw M, Konig R, Stertz S, Garcia-Sastre A, Chanda SK. Meta- and Orthogonal Integration of Influenza “OMICs” Data Defines a Role for UBR4 in Virus Budding. Cell Host Microbe. 2015;18(6):723–35. doi: 10.1016/j.chom.2015.11.002. PubMed PMID: 26651948; PMCID: PMC4829074.

58. Delorme G, Saltel F, Bonnelye E, Jurdic P, Machuca-Gayet I. Expression and function of semaphorin 7A in bone cells. Biol Cell. 2005;97(7):589–97. doi: 10.1042/BC20040103. PubMed PMID: 15859945.

59. Jaimes Y, Gras C, Goudeva L, Buchholz S, Eiz-Vesper B, Seltsam A, Immenschuh S, Blasczyk R, Figueiredo C. Semaphorin 7A inhibits platelet production from CD34+ progenitor cells. J Thromb Haemost. 2012;10(6):1100–8. doi: 10.1111/j.1538-7836.2012.04708.x. PubMed PMID: 22448926.

60. Ogata H, Goto S, Sato K, Fujibuchi W, Bono H, Kanehisa M. KEGG: Kyoto Encyclopedia of Genes and Genomes. Nucleic Acids Res. 1999;27(1):29–34. Epub 1998/12/10. doi: 10.1093/nar/27.1.29. PubMed PMID: 9847135; PMCID: PMC148090.

61. Liu X, Adorno-Cruz V, Chang YF, Jia Y, Kawaguchi M, Dashzeveg NK, Taftaf R, Ramos EK, Schuster EJ, El-Shennawy L, Patel D, Zhang Y, Cristofanilli M, Liu H. EGFR inhibition blocks cancer stem cell clustering and lung metastasis of triple negative breast cancer. Theranostics. 2021;11(13):6632–43. Epub 2021/05/18. doi: 10.7150/thno.57706. PubMed PMID: 33995681; PMCID: PMC8120216.

62. Lamiaa El-Shennawy ADH, Nurmaa K. Dashzeveg, Paul J. Mehl, Zihao Yu, Valerie L. Tokars, Vlad Nicolaescu, Carolina Ostiguin, Yuzhi Jia, Lin Li, Kevin Furlong, Chengsheng Mao, Jan Wysocki, Daniel Batlle, Thomas J. Hope, Yang Shen, Yuan Luo, Young Chae, Hui Zhang, Suchitra Swaminathan, Glenn C. Randall, Alexis R Demonbreun, Michael G Ison, Deyu Fang, Huiping Liu. Circulating ACE2-expressing Exosomes Block SARS-CoV-2 Infection as an Innate Antiviral Mechanism. bioRxiv. 2020. doi: https://doi.org/10.1101/2020.12.03.407031.

63. Kibria G, Ramos EK, Lee KE, Bedoyan S, Huang S, Samaeekia R, Athman JJ, Harding CV, Lotvall J, Harris L, Thompson CL, Liu H. A rapid, automated surface protein profiling of single circulating exosomes in human blood. Sci Rep. 2016;6:36502. doi: 10.1038/srep36502. PubMed PMID: 27819324; PMCID: PMC5098148.

64. Vences-Catalan F, Rajapaksa R, Srivastava MK, Marabelle A, Kuo CC, Levy R, Levy S. Tetraspanin CD81 promotes tumor growth and metastasis by modulating the functions of T regulatory and myeloid-derived suppressor cells. Cancer Res. 2015;75(21):4517–26. doi: 10.1158/0008-5472.CAN-15-1021. PubMed PMID: 26329536.

65. Maecker HT, Levy S. Normal lymphocyte development but delayed humoral immune response in CD81-null mice. J Exp Med. 1997;185(8):1505–10. PubMed PMID: 9126932; PMCID: PMC2196279.

66. Quast T, Eppler F, Semmling V, Schild C, Homsi Y, Levy S, Lang T, Kurts C, Kolanus W. CD81 is essential for the formation of membrane protrusions and regulates Rac1-activation in adhesion-dependent immune cell migration. Blood. 2011;118(7):1818–27. doi: 10.1182/blood-2010-12-326595. PubMed PMID: WOS:000294011500021.

67. van Zelm MC, Smet J, Adams B, Mascart F, Schandene L, Janssen F, Ferster A, Kuo CC, Levy S, van Dongen JJM, van der Burg M. CD81 gene defect in humans disrupts CD19 complex formation and leads to antibody deficiency. J Clin Invest. 2010;120(4):1265–74. doi: 10.1172/Jci39748. PubMed PMID: WOS:000276258100035.

68. Zoller M. Tetraspanins: push and pull in suppressing and promoting metastasis. Nat Rev Cancer. 2009;9(1):40–55. doi: 10.1038/nrc2543. PubMed PMID: 19078974.

69. Le Naour F, Andre M, Boucheix C, Rubinstein E. Membrane microdomains and proteomics: lessons from tetraspanin microdomains and comparison with lipid rafts. Proteomics. 2006;6(24):6447–54. doi: 10.1002/pmic.200600282. PubMed PMID: 17109380.

70. Zhu R, Gires O, Zhu L, Liu J, Li J, Yang H, Ju G, Huang J, Ge W, Chen Y, Lu Z, Wang H. TSPAN8 promotes cancer cell stemness via activation of sonic Hedgehog signaling. Nat Commun. 2019;10(1):2863. doi: 10.1038/s41467-019-10739-3. PubMed PMID: 31253779; PMCID: PMC6599078.

71. Podergajs N, Motaln H, Rajcevic U, Verbovsek U, Korsic M, Obad N, Espedal H, Vittori M, Herold-Mende C, Miletic H, Bjerkvig R, Turnsek TL. Transmembrane protein CD9 is glioblastoma biomarker, relevant for maintenance of glioblastoma stem cells. Oncotarget. 2016;7(1):593–609. doi: 10.18632/oncotarget.5477. PubMed PMID: 26573230; PMCID: PMC4808020.

72. Lee D, Na J, Ryu J, Kim HJ, Nam SH, Kang M, Jung JW, Lee MS, Song HE, Choi J, Lee GH, Kim TY, Chung JK, Park KH, Kim SH, Kim H, Seo H, Kim P, Youn H, Lee JW. Interaction of tetraspan(in) TM4SF5 with CD44 promotes self-renewal and circulating capacities of hepatocarcinoma cells. Hepatology. 2015;61(6):1978–97. doi: 10.1002/hep.27721. PubMed PMID: 25627085.

73. Bartee E, Eyster CA, Viswanathan K, Mansouri M, Donaldson JG, Fruh K. Membrane-Associated RING-CH proteins associate with Bap31 and target CD81 and CD44 to lysosomes. PLoS One. 2010;5(12):e15132. doi: 10.1371/journal.pone.0015132. PubMed PMID: 21151997; PMCID: PMC2996310.

74. Chen W, Patel D, Jia Y, Yu Z, Liu X, Shi H, Liu H. MARCH8 Suppresses Tumor Metastasis and Mediates Degradation of STAT3 and CD44 in Breast Cancer Cells. Cancers (Basel). 2021;13(11). Epub 2021/06/03. doi: 10.3390/cancers13112550. PubMed PMID: 34067416; PMCID: PMC8196951.

75. Giordano A, Gao H, Anfossi S, Cohen E, Mego M, Lee BN, Tin S, De Laurentiis M, Parker CA, Alvarez RH, Valero V, Ueno NT, De Placido S, Mani SA, Esteva FJ, Cristofanilli M, Reuben JM. Epithelial-mesenchymal transition and stem cell markers in patients with HER2-positive metastatic breast cancer. Mol Cancer Ther. 2012;11(11):2526–34. doi: 10.1158/1535-7163.MCT-12-0460. PubMed PMID: 22973057; PMCID: PMC3500676.

76. Ksiazkiewicz M, Markiewicz A, Zaczek AJ. Epithelial-mesenchymal transition: a hallmark in metastasis formation linking circulating tumor cells and cancer stem cells. Pathobiology. 2012;79(4):195–208. doi: 10.1159/000337106. PubMed PMID: 22488297.

77. Aktas B, Tewes M, Fehm T, Hauch S, Kimmig R, Kasimir-Bauer S. Stem cell and epithelial-mesenchymal transition markers are frequently overexpressed in circulating tumor cells of metastatic breast cancer patients. Breast Cancer Res. 2009;11(4):R46. doi: 10.1186/bcr2333. PubMed PMID: 19589136; PMCID: PMC2750105.

78. Dahan P, Martinez Gala J, Delmas C, Monferran S, Malric L, Zentkowski D, Lubrano V, Toulas C, Cohen-Jonathan Moyal E, Lemarie A. Ionizing radiations sustain glioblastoma cell dedifferentiation to a stem-like phenotype through survivin: possible involvement in radioresistance. Cell Death Dis. 2014;5:e1543. doi: 10.1038/cddis.2014.509. PubMed PMID: 25429620; PMCID: PMC4260760.

79. Dawood S, Austin L, Cristofanilli M. Cancer stem cells: implications for cancer therapy. Oncology (Williston Park). 2014;28(12):1101–7, 10. PubMed PMID: 25510809.

80. Giuliano M, Giordano A, Jackson S, Hess KR, De Giorgi U, Mego M, Handy BC, Ueno NT, Alvarez RH, De Laurentiis M, De Placido S, Valero V, Hortobagyi GN, Reuben JM, Cristofanilli M. Circulating tumor cells as prognostic and predictive markers in metastatic breast cancer patients receiving first-line systemic treatment. Breast Cancer Res. 2011;13(3):R67. doi: 10.1186/bcr2907. PubMed PMID: 21699723; PMCID: PMC3218956.

81. Pierga JY, Hajage D, Bachelot T, Delaloge S, Brain E, Campone M, Dieras V, Rolland E, Mignot L, Mathiot C, Bidard FC. High independent prognostic and predictive value of circulating tumor cells compared with serum tumor markers in a large prospective trial in first-line chemotherapy for metastatic breast cancer patients. Ann Oncol. 2012;23(3):618–24. doi: 10.1093/annonc/mdr263. PubMed PMID: 21642515.

82. Wang Z, Zhao K, Hackert T, Zoller M. CD44/CD44v6 a Reliable Companion in Cancer-Initiating Cell Maintenance and Tumor Progression. Front Cell Dev Biol. 2018;6:97. doi: 10.3389/fcell.2018.00097. PubMed PMID: 30211160; PMCID: PMC6122270.

83. Yu M, Bardia A, Wittner BS, Stott SL, Smas ME, Ting DT, Isakoff SJ, Ciciliano JC, Wells MN, Shah AM, Concannon KF, Donaldson MC, Sequist LV, Brachtel E, Sgroi D, Baselga J, Ramaswamy S, Toner M, Haber DA, Maheswaran S. Circulating breast tumor cells exhibit dynamic changes in epithelial and mesenchymal composition. Science. 2013;339(6119):580–4. doi: 10.1126/science.1228522. PubMed PMID: 23372014; PMCID: PMC3760262.

84. Uretmen Kagiali ZC, Sanal E, Karayel O, Polat AN, Saatci O, Ersan PG, Trappe K, Renard BY, Onder TT, Tuncbag N, Sahin O, Ozlu N. Systems-level Analysis Reveals Multiple Modulators of Epithelial-mesenchymal Transition and Identifies DNAJB4 and CD81 as Novel Metastasis Inducers in Breast Cancer. Mol Cell Proteomics. 2019;18(9):1756–71. doi: 10.1074/mcp.RA119.001446. PubMed PMID: 31221721; PMCID: PMC6731077.

85. Nagai J, Ishida Y, Koga N, Tanaka Y, Ohnuma K, Toyoda Y, Katoh A, Hayabuchi Y, Kigasawa H. A new sensitive and specific combination of CD81/CD56/CD45 monoclonal antibodies for detecting circulating neuroblastoma cells in peripheral blood using flow cytometry. J Pediatr Hematol Oncol. 2000;22(1):20–6. doi: 10.1097/00043426-200001000-00004. PubMed PMID: 10695817.

86. Al-Hajj M, Wicha MS, Benito-Hernandez A, Morrison SJ, Clarke MF. Prospective identification of tumorigenic breast cancer cells. P Natl Acad Sci USA. 2003;100(7):3983–8. doi: 10.1073/pnas.0530291100. PubMed PMID: WOS:000182058400082.

87. Dobin A, Davis CA, Schlesinger F, Drenkow J, Zaleski C, Jha S, Batut P, Chaisson M, Gingeras TR. STAR: ultrafast universal RNA-seq aligner. Bioinformatics. 2013;29(1):15–21. doi: 10.1093/bioinformatics/bts635. PubMed PMID: 23104886; PMCID: PMC3530905.

88. Love MI, Huber W, Anders S. Moderated estimation of fold change and dispersion for RNA-seq data with DESeq2. Genome Biol. 2014;15(12):550. doi: 10.1186/s13059-014-0550-8. PubMed PMID: 25516281; PMCID: PMC4302049.

89. Anders S, Pyl PT, Huber W. HTSeq--a Python framework to work with high-throughput sequencing data. Bioinformatics. 2015;31(2):166–9. doi: 10.1093/bioinformatics/btu638. PubMed PMID: 25260700; PMCID: PMC4287950.

90. Mertins P, Tang LC, Krug K, Clark DJ, Gritsenko MA, Chen L, Clauser KR, Clauss TR, Shah P, Gillette MA, Petyuk VA, Thomas SN, Mani DR, Mundt F, Moore RJ, Hu Y, Zhao R, Schnaubelt M, Keshishian H, Monroe ME, Zhang Z, Udeshi ND, Mani D, Davies SR, Townsend RR, Chan DW, Smith RD, Zhang H, Liu T, Carr SA. Reproducible workflow for multiplexed deep-scale proteome and phosphoproteome analysis of tumor tissues by liquid chromatography-mass spectrometry. Nat Protoc. 2018;13(7):1632–61. Epub 2018/07/11. doi: 10.1038/s41596-018-0006-9. PubMed PMID: 29988108; PMCID: PMC6211289.

91. Tsai CF, Zhao R, Williams SM, Moore RJ, Schultz K, Chrisler WB, Pasa-Tolic L, Rodland KD, Smith RD, Shi T, Zhu Y, Liu T. An Improved Boosting to Amplify Signal with Isobaric Labeling (iBASIL) Strategy for Precise Quantitative Single-cell Proteomics. Mol Cell Proteomics. 2020;19(5):828–38. Epub 2020/03/05. doi: 10.1074/mcp.RA119.001857. PubMed PMID: 32127492; PMCID: PMC7196584.

92. Okuda S, Watanabe Y, Moriya Y, Kawano S, Yamamoto T, Matsumoto M, Takami T, Kobayashi D, Araki N, Yoshizawa AC, Tabata T, Sugiyama N, Goto S, Ishihama Y. jPOSTrepo: an international standard data repository for proteomes. Nucleic Acids Res. 2017;45(D1):D1107–D11. Epub 2016/12/03. doi: 10.1093/nar/gkw1080. PubMed PMID: 27899654; PMCID: PMC5210561.

93. Vizcaino JA, Deutsch EW, Wang R, Csordas A, Reisinger F, Rios D, Dianes JA, Sun Z, Farrah T, Bandeira N, Binz PA, Xenarios I, Eisenacher M, Mayer G, Gatto L, Campos A, Chalkley RJ, Kraus HJ, Albar JP, Martinez-Bartolome S, Apweiler R, Omenn GS, Martens L, Jones AR, Hermjakob H. ProteomeXchange provides globally coordinated proteomics data submission and dissemination. Nat Biotechnol. 2014;32(3):223–6. Epub 2014/04/15. doi: 10.1038/nbt.2839. PubMed PMID: 24727771; PMCID: PMC3986813.

94. Kong AT, Leprevost FV, Avtonomov DM, Mellacheruvu D, Nesvizhskii AI. MSFragger: ultrafast and comprehensive peptide identification in mass spectrometry-based proteomics. Nat Methods. 2017;14(5):513–20. Epub 2017/04/11. doi: 10.1038/nmeth.4256. PubMed PMID: 28394336; PMCID: PMC5409104.

95. Teo GC, Polasky DA, Yu F, Nesvizhskii AI. Fast Deisotoping Algorithm and Its Implementation in the MSFragger Search Engine. J Proteome Res. 2021;20(1):498–505. Epub 2020/12/18. doi: 10.1021/acs.jproteome.0c00544. PubMed PMID: 33332123.

96. Tyanova S, Temu T, Sinitcyn P, Carlson A, Hein MY, Geiger T, Mann M, Cox J. The Perseus computational platform for comprehensive analysis of (prote)omics data. Nat Methods. 2016;13(9):731–40. Epub 2016/06/28. doi: 10.1038/nmeth.3901. PubMed PMID: 27348712.

97. Jiao X, Sherman BT, Huang da W, Stephens R, Baseler MW, Lane HC, Lempicki RA. DAVID-WS: a stateful web service to facilitate gene/protein list analysis. Bioinformatics. 2012;28(13):1805–6. Epub 2012/05/01. doi: 10.1093/bioinformatics/bts251. PubMed PMID: 22543366; PMCID: PMC3381967.

98. Szklarczyk D, Gable AL, Nastou KC, Lyon D, Kirsch R, Pyysalo S, Doncheva NT, Legeay M, Fang T, Bork P, Jensen LJ, von Mering C. The STRING database in 2021: customizable protein-protein networks, and functional characterization of user-uploaded gene/measurement sets. Nucleic Acids Res. 2021;49(D1):D605–D12. Epub 2020/11/26. doi: 10.1093/nar/gkaa1074. PubMed PMID: 33237311; PMCID: PMC7779004.

99. Shannon P, Markiel A, Ozier O, Baliga NS, Wang JT, Ramage D, Amin N, Schwikowski B, Ideker T. Cytoscape: a software environment for integrated models of biomolecular interaction networks. Genome Res. 2003;13(11):2498–504. Epub 2003/11/05. doi: 10.1101/gr.1239303. PubMed PMID: 14597658; PMCID: PMC403769.

